# Biomimetic MRI Nanoprobe for Mapping Cerebrovascular Inflammation After Traumatic Brain Injury

**DOI:** 10.64898/2026.05.25.727244

**Authors:** Joshua Rousseau, Amberlyn Simmons, Michelle Kim, Dakota Ortega, Owen Alzubi, Ting-Yun Wang, Benjamin B. Bartelle, Sarah E. Stabenfeldt, Kuei-Chun Wang

**Affiliations:** School of Biological and Health Systems Engineering, Arizona State University, Tempe, AZ, USA; School of Computing and Augmented Intelligence, Arizona State University, Tempe, AZ, USA; Knowledge Enterprise Biosciences Research Preclinical Imaging Center, Arizona State University, Tempe, AZ, USA; John Shufeldt School of Medicine and Medical Engineering, Arizona State University, Phoenix, AZ, USA

## Abstract

Cerebrovascular inflammation is a critical driver of secondary injury following traumatic brain injury (TBI), yet no noninvasive tool currently exists to map its spatial heterogeneity across the injured brain. Here, we report MoNP-SPION, a monocyte-mimetic nanoprobe that selectively targets activated cerebrovascular endothelium after TBI and enables quantitative MRI of spatiotemporally heterogeneous cerebrovascular inflammation. MoNP-SPION binding to inflamed brain endothelium scaled with activation state, enabling rapid, spatially heterogeneous targeting of injured cerebral vasculature. MoNP-SPION-enhanced MRI revealed dynamic reductions in T2* throughout the brain that extended beyond the injury penumbra and captured both subacute vascular injury and therapy-induced recovery. Importantly, spatiotemporal reductions in T2* strongly correlated with VCAM1 upregulation, supporting MoNP-SPION as a molecularly specific readout of endothelial activation. Together, these findings establish MoNP-SPION as a promising platform for longitudinal, noninvasive monitoring of cerebrovascular inflammation in TBI and other neurological disorders involving vascular dysfunction.

**TEASER:** Cell-mimicking MRI nanoprobe maps cerebrovascular injury and recovery dynamics following brain trauma.

## 1. INTRODUCTION

Traumatic brain injury (TBI), resulting from sudden mechanical impact to the head, remains a leading cause of mortality and long-term disability worldwide (*1*). The primary insult causes immediate and irreversible structural damage to the brain, including contusions, lacerations, and intracranial hemorrhage, and trigger a cascade of secondary pathological processes that drive neurological decline, including cerebrovascular inflammation (*2*). Immediately after injury, mechanical stimuli together with cytokines and chemokines released by activated microglia and astrocytes shift endothelial cells (EC) lining the cerebral vasculature into a pro-inflammatory phenotype. This inflammation is characterized by upregulation of cell adhesion molecules, including vascular cell adhesion molecule 1 (VCAM1) and intercellular adhesion molecule 1 (ICAM1), and the disruption of intercellular junctions, compromising blood-brain barrier (BBB) integrity (*3*, *4*). This acute inflammation serves a critical role in initiating wound healing in the brain, yet chronic vascular inflammation may promote continued leukocyte infiltration into the brain parenchyma, thereby amplifying neuroinflammation and contributing to progressive tissue damage (*5*, *6*). Importantly, post-TBI cerebrovascular inflammation is neither spatially confined to the primary lesion nor transient but extends into distinct cortical and subcortical regions and evolves over time (*7*). When unresolved, chronic cerebrovascular inflammation leads to progressive neurodegeneration and vascular cognitive impairment (VCI) (*8*).

Despite its clinical significance, noninvasive tools for detecting cerebrovascular inflammation with spatiotemporal precision after TBI remain lacking (*9*). Current diagnostic strategies for TBI rely primarily on computed tomography (CT), conventional magnetic resonance imaging (MRI), and clinical scoring systems such as the Glasgow Coma Scale (GCS). While these approaches are valuable for identifying structural damage, such as hemorrhage or edema, and assessing neurological impairment, they provide little information about the underlying cerebrovascular inflammation that drives secondary injury (*10*). Advanced MRI techniques, including dynamic contrast-enhanced (DCE) MRI and diffusion-weighted imaging (DWI), offer improved sensitivity to BBB disruption and axonal injury in patients with TBI, respectively (*11*, *12*). Yet these methods remain indirect and are insensitive to the cellular and molecular features of cerebrovascular inflammation. Molecular imaging strategies such as translocator protein positron emission tomography (TSPO-PET) have been investigated to address this limitation (*13*). However, TSPO-PET primarily reflects glial activation rather than inflammatory burden within the cerebral vasculature and is further constrained by the limited spatial resolution and suboptimal radioligand specificity. Collectively, these gaps highlight the need for molecular imaging tools capable of directly mapping cerebrovascular inflammation following TBI.

Iron oxide nanoparticles have been extensively explored as MRI contrast agents given their strong T2/T2* relaxivity. Ultrasmall superparamagnetic iron oxide nanoparticles (SPIONs), in particular, can extravasate through regions of compromised BBB integrity, extending their utility beyond reticuloendothelial imaging of the liver and spleen to the detection of neuroinflammatory pathologies (*14*). However, their translational potential remains constrained by nonspecific and inefficient accumulation, which often necessitates delayed imaging protocols and limits their suitability for acute injury settings (*15*). The selective upregulation of surface adhesion molecules on activated endothelium provides a molecularly distinct opportunity to enhance contrast agent accumulation at sites of cerebrovascular inflammation. Indeed, conjugating VCAM1- or ICAM1-specific antibodies to SPION surfaces have demonstrated improved vascular selectivity in preclinical stroke and neuroinflammation models (*16*, *17*). Beyond ligand-based targeting approaches, biomimetic nanoparticle platforms offer a distinct strategy for enhancing site-specific SPION delivery to vascular inflammation. We recently engineered monocyte membrane-coated, SPION-encapsulated polymeric nanoparticles (MoNP-SPION) that leverage the multivalent, receptor-diverse characteristics of native leukocyte-endothelial interactions. MoNP-SPION demonstrated preferential accumulation at sites of vascular inflammation and enhanced MRI detection of plaques in mouse models of atherosclerosis (*18*). Here, we extend this platform to TBI, where the spatiotemporal dynamics of cerebrovascular inflammation remain incompletely defined and molecular imaging strategies capable of distinguishing endothelial activation from nonspecific barrier disruption are critically needed.

In this study, we evaluated MoNP-SPION as a molecular MRI nanoprobe for imaging cerebrovascular inflammation after TBI. We first demonstrate that monocyte membrane coating significantly enhances nanoparticle localization to the injured brain in a controlled cortical impact (CCI) mouse model, where fluorescent MoNP colocalize with VCAM1-positive endothelium throughout the injured cerebral vasculature via β1 integrin-dependent interactions. Building on this targeting specificity, we further show that MoNP-SPION generate spatially heterogeneous T2* contrast enhancement in the injured brain distinct from uncoated counterparts, and that longitudinal imaging captures the spatiotemporal evolution of cerebrovascular inflammation with regional signal patterns supported by VCAM1 staining. Moreover, pharmacological suppression of inflammation with minocycline (Mc) or atorvastatin (At) correspondingly attenuates MoNP-SPION contrast *in vivo* to varying degrees. Together, these findings establish MoNP-SPION-enhanced MRI as a sensitive and biologically specific platform for imaging cerebrovascular inflammation after TBI, with potential utility for monitoring neuroinflammatory progression and therapeutic response.

## 2. RESULTS

### 2.1 Fluorescent MoNP selectively bind inflamed brain ECs and TBI vasculature

The brain microvasculature represents a uniquely challenging vascular bed, where the intact BBB restricts nanoparticle access and endothelial activation patterns differ from those of peripheral vessels (*19*). While our MoNP platform has previously been shown to target inflamed arterial and venous ECs, its ability to engage inflamed brain microvascular endothelium remains unknown (*18*, *20*). Establishing this capability is essential for extending MoNP-based targeting to neurovascular injury and disease. To address this, we first prepared fluorescently labeled MoNP and confirmed that their physicochemical properties were consistent with our previously reported MoNP formulations (**Fig. S1**) (*20*). We then investigated MoNP interactions with human brain microvascular ECs (HBMECs) *in vitro* and in a murine CCI model (**Fig. 1A**). Confocal microscopy revealed that MoNP were effectively internalized by tumor necrosis factor α (TNFα)-activated HBMECs, whereas uncoated polymeric nanoparticle cores (NP) showed minimal uptake (**Fig. 1B**). We next asked whether MoNP-HBMEC interactions scale with the degree of endothelial activation. HBMECs were treated with increasing concentrations of TNFα to induce graded inflammatory activation. Western blot analysis demonstrated that VCAM1 expression increased with TNFα concentration and plateaued at 10 ng/mL (**Fig. S2**). Using selected inflammatory conditions (0, 0.1, 1, and 10 ng/mL of TNFα), HBMECs were incubated with MoNP. Quantification of fluorescent intensity revealed progressively greater MoNP uptake with increasing endothelial activation state (**Fig. 1C**). This trend was further confirmed by flow cytometry, demonstrating inflammation-dependent increases in intracellular MoNP fluorescence (**Fig. 1D**). To evaluate the extent of accumulation *in vivo* during heightened cerebrovascular inflammation, MoNP were administered via retro-orbital injection (RO) to naïve control mice or inured mice 24 hours post-CCI. *In vivo* imaging system (IVIS) analysis of excised brains showed no detectable fluorescence in all naïve mice, while injured mice receiving uncoated NP exhibited only moderate signal localized to the injury site (**Figs. 1E & S3A**). In contrast, injured mice administered MoNP displayed significantly stronger fluorescence at the injury core, with signal extending into adjacent brain regions and decreasing with distance from the injury. Immunofluorescence (IF) analysis of brain sections showed that, while injured NP-treated mice exhibited minimal DiD signal, MoNP colocalized with VCAM1- and CD31-positive cells, indicating direct interaction with inflamed endothelium (**Figs. 1F & S3B**).

**Figure 1.**
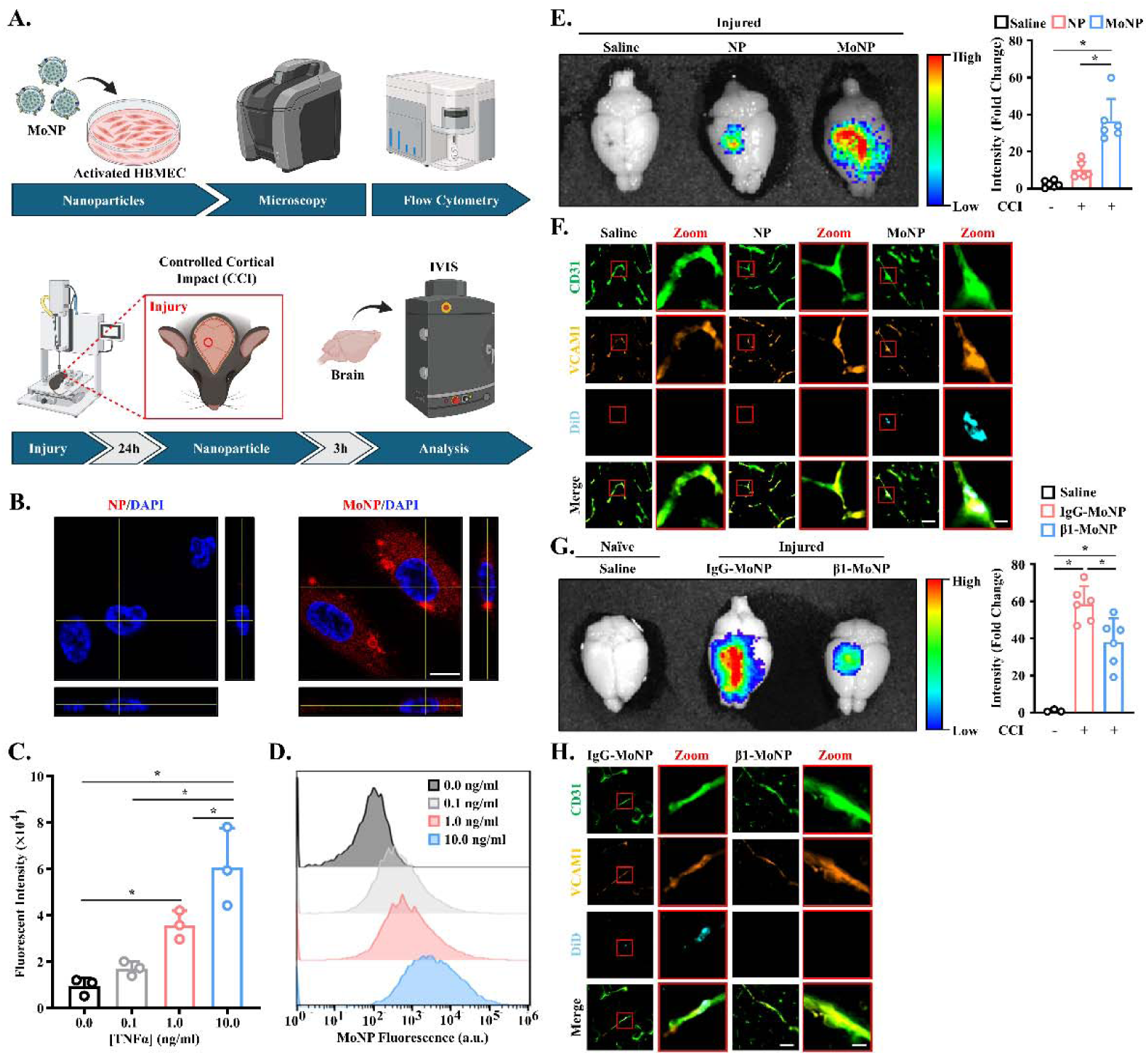
Fluorescent MoNP selectively bind inflamed brain ECs and accumulate in CCI-injured brain vasculature. (A) Schematic diagram of *in vitro* and *in vivo* experiments. (B) Representative confocal microscope images of TNFα-activated HBMECs treated with MoNP and NP; scale bar = 15 µm. (C) Quantification of MoNP uptake to varying degrees of TNFα-activated HBMECs; n = 3. (D) Flow cytometric analysis of MoNP uptake to varying degrees of TNFα-activated HBMECs; n = 3. (E) Representative IVIS images and whole brain fluorescent signal quantification of injured mouse brains after administration of MoNP, NP, or saline; n = 6. (F) Representative IF images of DiD colocalization with CD31 and VCAM1 expression in the ipsilateral hemisphere; scale bar (left) = 50 µm, scale bar (right) = 10 µm. (G) Representative IVIS images and whole brain fluorescent signal quantification of naïve and injured mouse brains after administration of saline, IgG-MoNP, or β1-MoNP; n = 3 for naïve saline, n = 6 for all others. (H) Representative IF images of DiD colocalization with CD31 and VCAM1 expression in the ipsilateral hemisphere; scale bar (left) = 50 µm, scale bar (right) = 10 µm. All data is presented as mean ± SD. * indicates p < 0.05. Statistical significance was calculated with one-way ANOVA with Tukey’s post-hoc multiple comparisons test.

Monocytes engage activated endothelium through very late antigen 4 (VLA4; α4β1 integrin) (*21*). To determine whether MoNP exploit the same mechanisms to interact with inflamed brain endothelium, MoNP pretreated with β1-integrin blocking antibody (β1-MoNP) or IgG isotype control (IgG-MoNP) were administered to injured mice. IgG-MoNP maintained widespread distribution surrounding the injury, whereas β1-MoNP were largely restricted to the injury core (**Fig. 1G**). Consistently, IF demonstrated colocalization of IgG-MoNP with inflamed brain ECs, while β1-MoNP showed minimal endothelial association (**Fig. 1H**). Collectively, these findings indicate that MoNP binding to brain endothelium depends on local inflammatory cues and mediated through β1-integrin–mediated interactions, supporting the use of this platform for detecting regional differences in cerebrovascular inflammation following brain injury.

### 2.2 MoNP-SPION generates dynamic MRI contrast *in vitro* and in TBI mice

To evaluate the MR imaging capabilities of MoNP in TBI-induced cerebrovascular inflammation, we generated MoNP-SPION by incorporating SPIONs into the MoNP platform via a modified double emulsion method. Dynamic light scattering (DLS) and transmission electron microscopy (TEM) analyses confirmed that MoNP-SPION maintained physicochemical properties, including a hydrodynamic diameter of ∼270 nm and low polydispersity, consistent with our previously reported formulation (**Figs. S4**) (*18*). We next incubated TNFα-activated HBMECs with MoNP-SPION and assessed nanoparticle uptake using Prussian Blue staining and MR imaging (**Fig. 2A**). Microscopy analysis of Prussian Blue-stained cells confirmed the preferential MoNP-SPION accumulation in TNFα-activated HBMECs (**Fig. 2B**). In HBMECs treated with increasing TNFα concentrations, MoNP-SPION generated progressively stronger hypointense contrast in cell phantoms (**Fig. 2C**). Cellular iron content, measured by a Prussian Blue-based colorimetric assay, increased proportionally with inflammatory stimulation, confirming that MR contrast changes reflected increased MoNP-SPION uptake (**Fig. 2D**). To determine the optimal *in vivo* imaging window, MoNP-SPION was administered via RO injection 24 hours post-CCI, followed by serial T2*-weighted MR imaging for the subsequent 24 hours. Imaging was performed using a thick (1 mm) imaging slice to enhance sensitivity for whole-region contrast quantification and temporal kinetic analysis. Rapid accumulation and hypointense contrast were observed within the injury penumbra, with peak contrast observed at 3 hours post-injection (**Fig. 2E**). Whole-brain, multi-echo T2* decay analysis similarly revealed maximal contrast differentiation at 3 hours after MoNP-SPION administration, whereas other timepoints were less distinct or more closely resembled baseline values (**Fig. 2F**). Quantification of T2* decay constants across the contralateral hemisphere, ipsilateral hemisphere, and injury penumbra demonstrated the lowest values at 3 hours, with progressive recovery toward baseline thereafter (**Fig. 2G**). Together, these results demonstrate that MoNP-SPION produces inflammation-dependent uptake *in vitro* and rapid, dynamic contrast changes *in vivo* with a 3-hour imaging window post-injection and efficient clearance thereafter.

**Figure 2.**
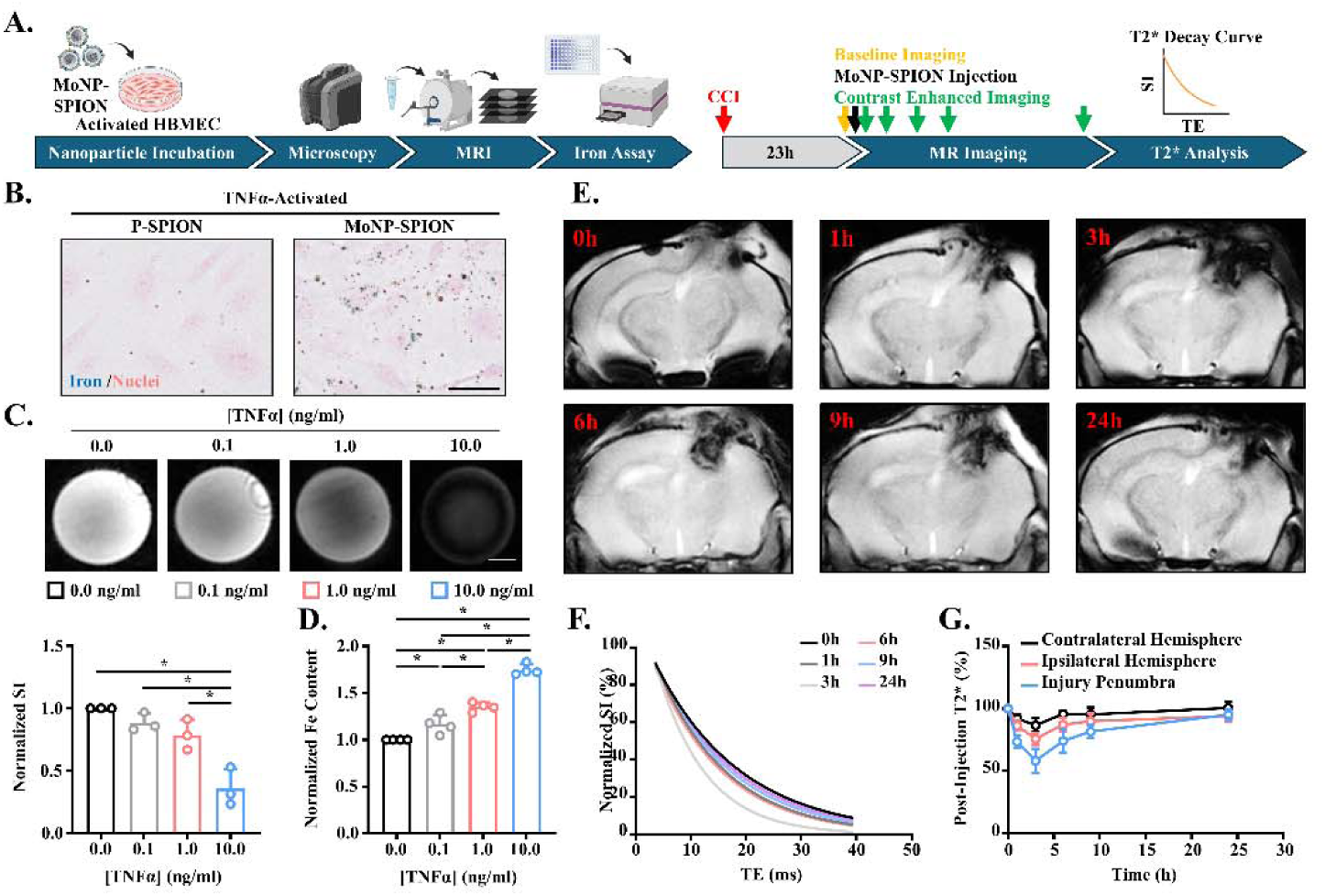
MoNP-SPION generates dynamic MRI contrast *in vitro* and in TBI mice. (A) Schematic diagram of *in vitro* and *in vivo* experimental design and workflow. (B) Representative Prussian Blue stained images of TNFα-activated HBMECs treated with MoNP-SPION and P-SPION; scale bar = 100 µm. (C) Representative T2*-weighted MRI phantom images and SI quantification of MoNP-SPION uptake to varying degrees of activated HBMECs; scale bar = 250 µm, n = 3. (D) Aqueous iron assay measurements of MoNP-SPION uptake to varying degrees of activated HBMECs; n = 4. (E) Representative coronal plane T2*-weighted MR images of TBI mice before and 1, 3, 6, 9, and 24 hours post-injection of MoNP-SPION. (F) Quantification of whole brain T2* decay curves before and 1, 3, 6, 9, and 24 hours post-injection of MoNP-SPION. (G) T2* decay constants represented as a percentage of baseline values over 24 hours within the contralateral hemisphere, ipsilateral hemisphere, and injury penumbra; n = 4. All data is presented as mean ± SD. * indicates p < 0.05. Statistical significance was calculated with one-way ANOVA with Tukey’s post-hoc multiple comparisons test.

### 2.3 MoNP-SPION produces spatially heterogeneous MRI contrast after TBI

Having established the optimal imaging window, we next employed a higher-resolution imaging strategy using 0.5-mm slices to minimize partial volume effects and resolve the spatial distribution of MoNP-SPION within the injured brain compared to non-targeted SPION formulation. Twenty-three hours after CCI injury, mice underwent baseline MRI imaging. At 24 hours post-CCI, MoNP-SPION, its uncoated counterpart P-SPION, or saline, were administered via RO injection, followed by MR imaging to compare the spatial contrast profiles 3 hours post-injection (**Fig. 3A**). Naïve mice, regardless of treatment, exhibited negligible contrast changes in T2* decay (**Fig. S5**). Among injured mice, administration of saline produced no appreciable contrast, whereas P-SPION generated mild hypointense signal largely confined to the injury penumbra (**Fig. 3B**). In contrast, MoNP-SPION produced robust hypointense speckling throughout the brain, accompanied by spatially heterogenous contrast and alterations in T2* relaxation (**Figs. 3B & 3C**). Quantification T2* decay analysis confirmed that P-SPION reduced T2* primarily within the penumbra, whereas MoNP-SPION significantly decreased T2* across both hemispheres as well as the injury region (**Fig. 3D**, **Table 1**). Consistent with the *in vivo* findings, *ex vivo* brain imaging further showed that MoNP-SPION retained a persistent speckled contrast pattern suggestive of vascular binding (**Fig. 3E**). Regional T2* analysis revealed the greatest reductions in areas nearest the injury, including the ipsilateral isocortex (IC) and ipsilateral diencephalon (DE), with progressively smaller reductions in more distal regions such as contralateral olfactory area and cortical subplate (OA/CS). In contrast, P-SPION produced only mild T2* reductions near the injury and minimal changes elsewhere, while saline treatments and all naïve brains produce no reduction in T2* (**Figs. 3F & S6, Table S1**). To further evaluate spatial distribution, multi-slice MR imaging analysis was performed on the *in vivo* dataset across rostral and caudal planes relative to the injury. The speckled MoNP-SPION contrast pattern persisted across planes spanning ±2.0 mm but decreased in intensity with distance from the injury site (**Fig. 3G & S7**). Quantification confirmed that while saline produced no measurable T2* alterations throughout the brain, both P-SPION and MoNP-SPION exhibited their lowest T2* decay values at the injury plane with progressive recovery distally; however, MoNP-SPION consistently produced greater contrast changes across all regions examined (**Figs. 3H & S8**). Prussian Blue staining confirmed the presence of iron deposits within the distinct brain regions of notable T2* contrast (**Fig. S9**). Collectively, these findings demonstrate that MoNP-SPION generates spatially heterogeneous contrast both within and beyond the injury penumbra, enabling three-dimensional mapping of cerebrovascular inflammation acutely following TBI.

**Figure 3.**
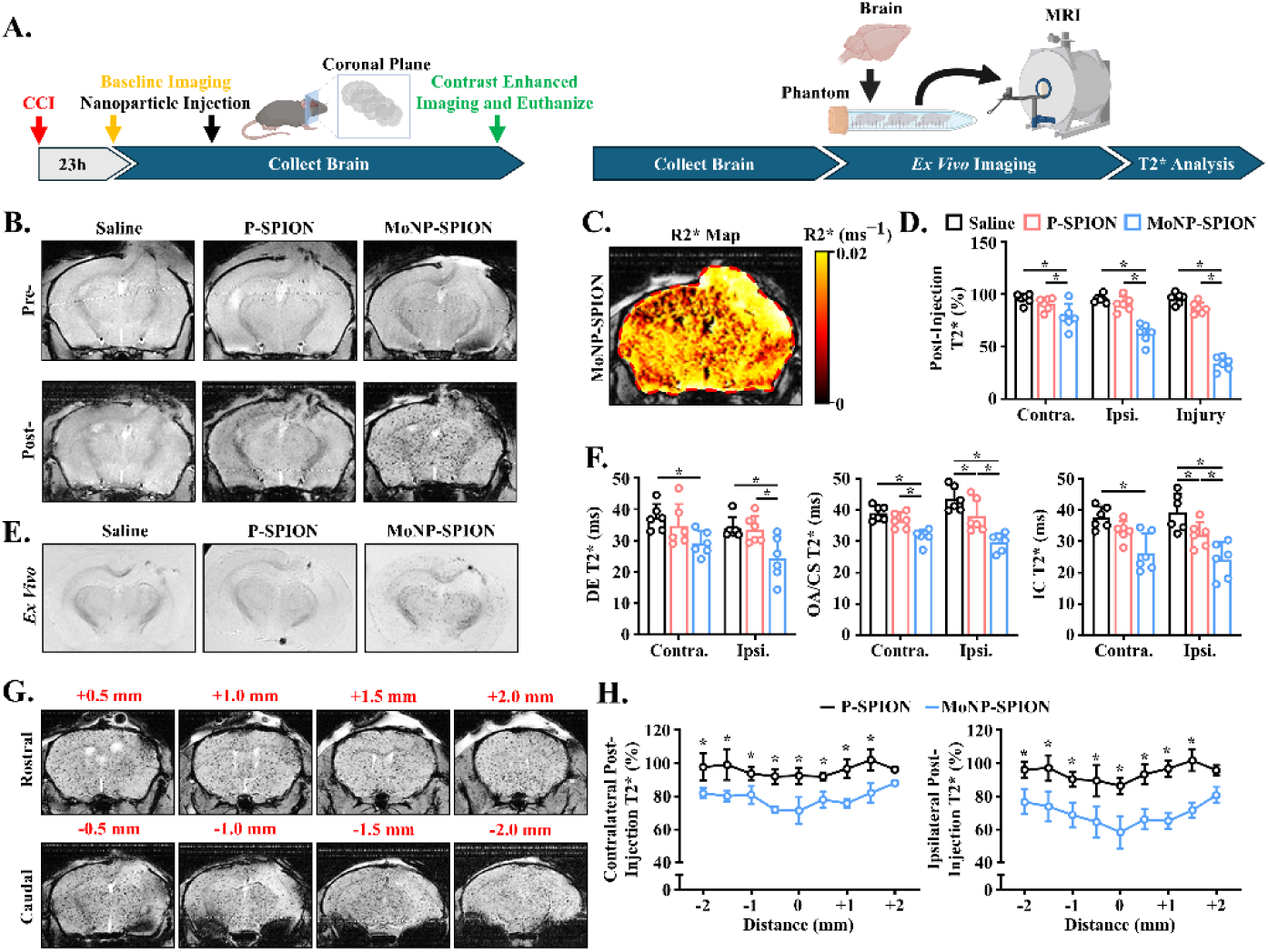
MoNP-SPION generates spatially heterogeneous MRI contrast after TBI. (A) Schematic diagram of MR imaging timeline. (B) Representative coronal plane T2*-weighted MR images of 1-day injured mouse brains before and 3 hours after administration of MoNP-SPION, P-SPION, or saline. (C) Representative image of R2* heat map. (D) Post-injection T2* values represented as a percentage of pre-injection values; n = 6. (E) Representative T2*-weighted *ex vivo* brain MR images of injured mice administered MoNP-SPION, P-SPION, or saline. (F) T2* quantification of the DE (left), OA/CS (center), and IC (right) within the contralateral and ipsilateral hemispheres; n = 6. (G) Representative multi-slice coronal plane T2*-weighted images of injured mouse brains after administration of MoNP-SPION from +2 mm rostral to -2 mm caudal of the injury core. (H) T2* quantification of brain sections +2 mm rostral to -2 mm caudal of the injury core within the contralateral (left) and ipsilateral (right) hemispheres represented as a percentage of baseline values; n = 3. All data is presented as mean ± SD, * indicates p < 0.05. * indicates p < 0.05. Statistical significance was calculated with one-way ANOVA with Tukey’s post-hoc multiple comparisons test.

**Table 1.**
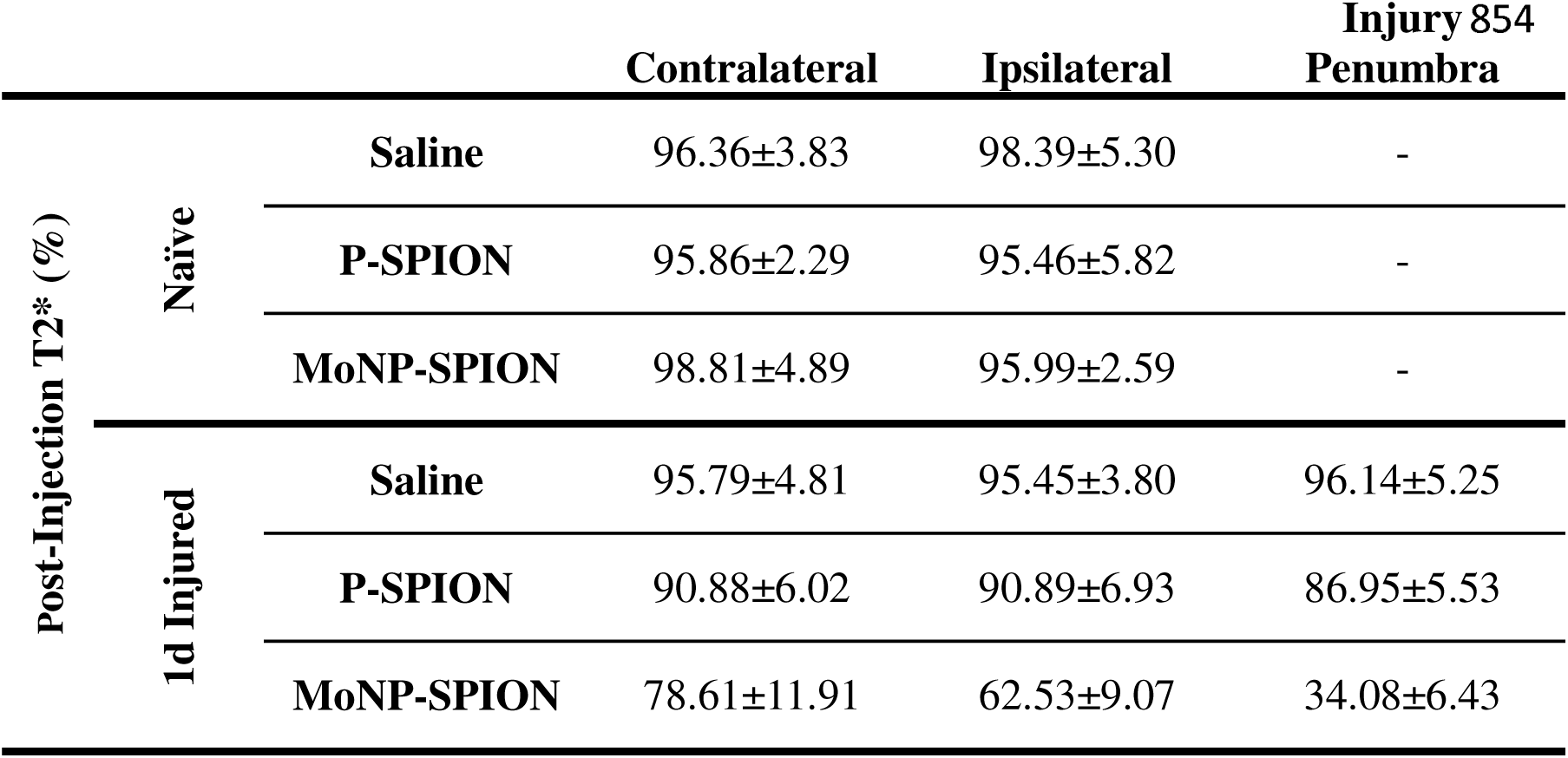
*In vivo* T2* analysis 1 day post-injury. Post-injection T2* decay constants represented as a percentage of pre-injection values across the contralateral hemisphere, ipsilateral hemisphere, and injury penumbra from mouse brains after administration of MoNP-SPION. P-SPION, or saline; n = 6. Data is represented as mean ± SD.

### 2.4 MoNP-SPION-Enhanced MRI reveals spatiotemporal evolution of contrast after TBI

The spatial heterogeneity of hypointense contrast observed at 24 hours post-injury prompted us to test whether MoNP-SPION could probe temporal evolution of cerebrovascular inflammation. We and others have shown that by 7-days post-injury the cerebrovascular inflammation is diminished relative to the 1-day post-injury response, as evidenced by reduced inflammatory cytokine levels and leukocyte recruitment (*22*). Therefore, we assessed the functionality of the MoNP-SPIONs to capture this temporal inflammatory dynamic where mice underwent CCI followed by MoNP-SPION–enhanced MR imaging at 7 days post-TBI, with naïve mice serving as controls. The pronounced speckled hypointense pattern observed at 1 day remained detectable at 7 days but was less prominent (**Fig. 4A**). Notably, the injury penumbra retained the strongest contrast at both time points, indicating that the spatial gradient of MoNP-SPION accumulation persisted into the subacute phase. Relaxivity maps supported these observations, exhibiting the greatest degree of spatial heterogeneity across the brain in 1-day injured mice compared to 7-day and naïve groups (**Fig. 4B**). T2* decay analysis across both hemispheres and the injury penumbra likewise showed partial recovery by day 7 relative to day 1, although values remained below naïve mice levels (**Fig. 4C**, **Table 2**). Multi-slice imaging along the z-axis further demonstrated persistent spatial heterogeneity, with contrast reductions observed across both hemispheres and diminishing with distance from the injury plane (**Fig. 4D**). *Ex viv*o MR imaging confirmed these *in vivo* observations, revealing retained but reduced speckled contrast throughout the brain at 7 days (**Fig. 4E**). Regional T2* decay quantification showed modest recovery across most brain regions, with minimal residual changes except in the contralateral OA/CS, which exhibited a persistent decrease at 7 days post-injury **(Fig. 4F, Table S2**). Collectively, these findings demonstrate that MoNP-SPION enables longitudinal mapping of the spatiotemporal evolution of cerebrovascular inflammation after TBI.

**Figure 4.**
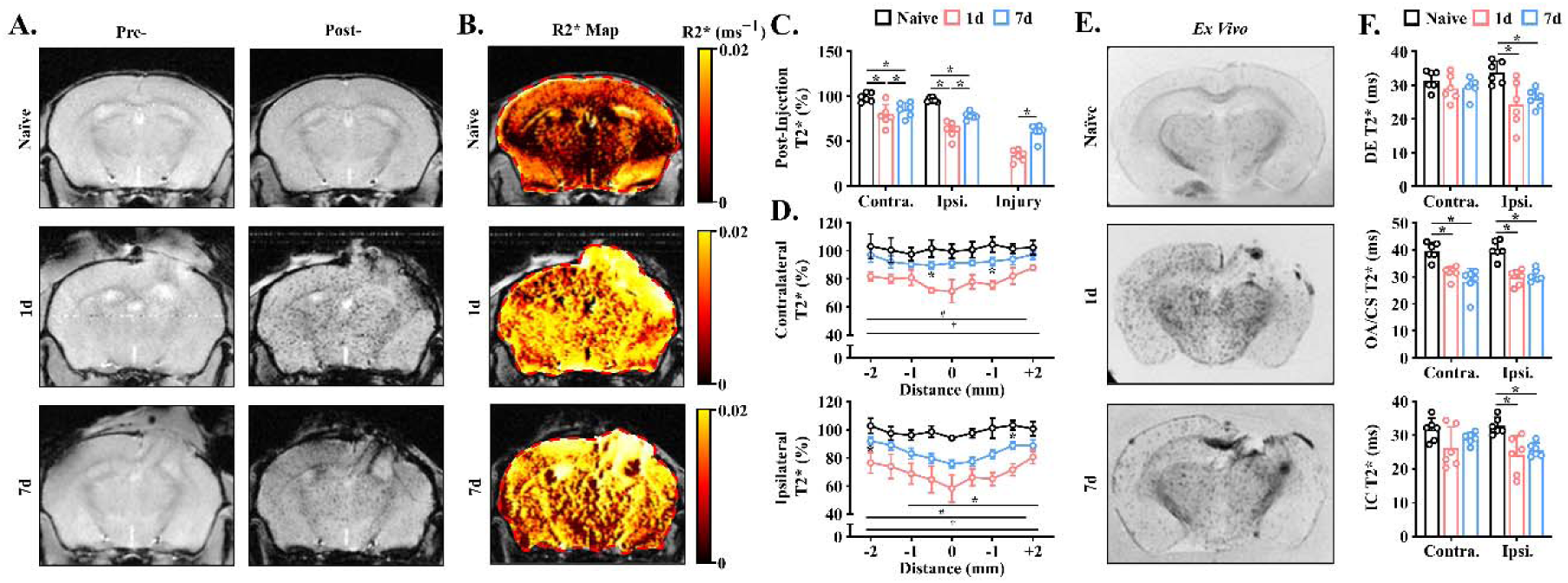
MoNP-SPION MRI reveals spatiotemporal evolution of brain contrast after TBI. (A) Representative T2*-weighted images of naïve, 1-day injured, and 7-day injured mouse brains before and 3 hours after administration of MoNP-SPION. (B) *In vivo* coronal plane R2* map of naïve, 1-day injured, and 7-day injured mouse brains. (C) T2* quantification of the contralateral hemisphere, ipsilateral hemisphere, and injury penumbra as a percentage of baseline values; n = 6. (D) T2* quantification of brain sections +2 mm rostral to -2 mm caudal of the injury core within the contralateral (top) and ipsilateral (bottom) hemispheres represented as a percentage of baseline values; n = 3, *p<0.05, naïve vs 7 day; #p<0.05, 1 day vs. 7 day; † p<0.05, naïve vs. 7 day. (E) Representative T2*-weighted *ex vivo* brain MR images of naïve, 1-day injured, and 7-day injured mice after administration of MoNP-SPION. (F) T2* quantification of the DE (top), OA/CS (middle), and IC (bottom) within the contralateral and ipsilateral hemispheres; n = 6. All data is presented as mean ± SD. *, #, and † indicate p < 0.05. Statistical significance was calculated with two-way ANOVA with Tukey’s post-hoc multiple comparisons test.

**Table 2:** *In vivo* spatiotemporal T2* analysis. Post-injection T2* decay constants represented as a percentage of pre-injection values across the contralateral hemisphere, ipsilateral hemisphere, and injury penumbra from mouse brains after administration of MoNP-SPION in naïve, 1 day injured, and 7 day injured mice; n = 6. Data is represented as mean ± SD.

| Post-Injection<br>T2* (%) |  | Contralateral | Ipsilateral | Injury 861<br>Penumbra |
| --- | --- | --- | --- | --- |
|  | Naïve | 98.81±4.89 | 95.99±2.59 | - |
|  | 1d Injured | 78.61±11.91 | 62.53±9.07 | 34.08±6.43 |
|  | 7d Injured | 85.18±7.94 | 78.20±4.13 | 60.44±8.41 |

### 2.5 MoNP-SPION contrast reflects regional endothelial activation after TBI

The preceding experiments established that MoNP-SPION produces spatially and temporally heterogeneous T2* contrast in the injured brain, but whether this signal directly reflects regional endothelial activation remains unclear. To address this, brain sections from naïve and CCI-injured mice were stained for VCAM1 to assess the spatiotemporal distribution of cerebrovascular inflammation. Macroscopic IF imaging revealed minimal VCAM1 expression in naïve brains, whereas 1-day post-injury brains exhibited marked VCAM1 upregulation concentrated near the injury penumbra. At 7 days post-injury, VCAM1 expression was reduced but remained detectable (**Fig. 5A**). Regional high-magnification imaging further supported this temporal pattern across the DE, OA/CS, and IC (**Fig. S10**). Focusing on the IC, VCAM1 expression in the contralateral hemisphere was mild at 1 day and slightly slower at 7 days, whereas the ipsilateral IC displayed robust VCAM1 expression at 1 day post-injury, which declined by day 7 but remained elevated relative to the contralateral hemisphere (**Fig. 5B**). Quantification of VCAM1-positive ECs confirmed these observations, with the ipsilateral IC at 1 day post-injury showing the highest percentage of VCAM1-positive cells and progressively lower expression in more distal regions. By day 7, VCAM1-positive ECs decreased across most brain regions to approximately 5%, except in the contralateral OA/CS, which showed a modest increase (**Fig. 5C**). Comparison of this regional VCAM1 expression with corresponding T2* decay revealed a strong negative correlation (Pearson’s coefficient = -0.7530, p = 0.0003) (**Fig. 5D**). Collectively, these findings support MoNP-SPION contrast as a molecular imaging readout associated with endothelial VCAM1 expression, thereby enabling spatiotemporal mapping of cerebrovascular inflammation.

**Figure 5.**
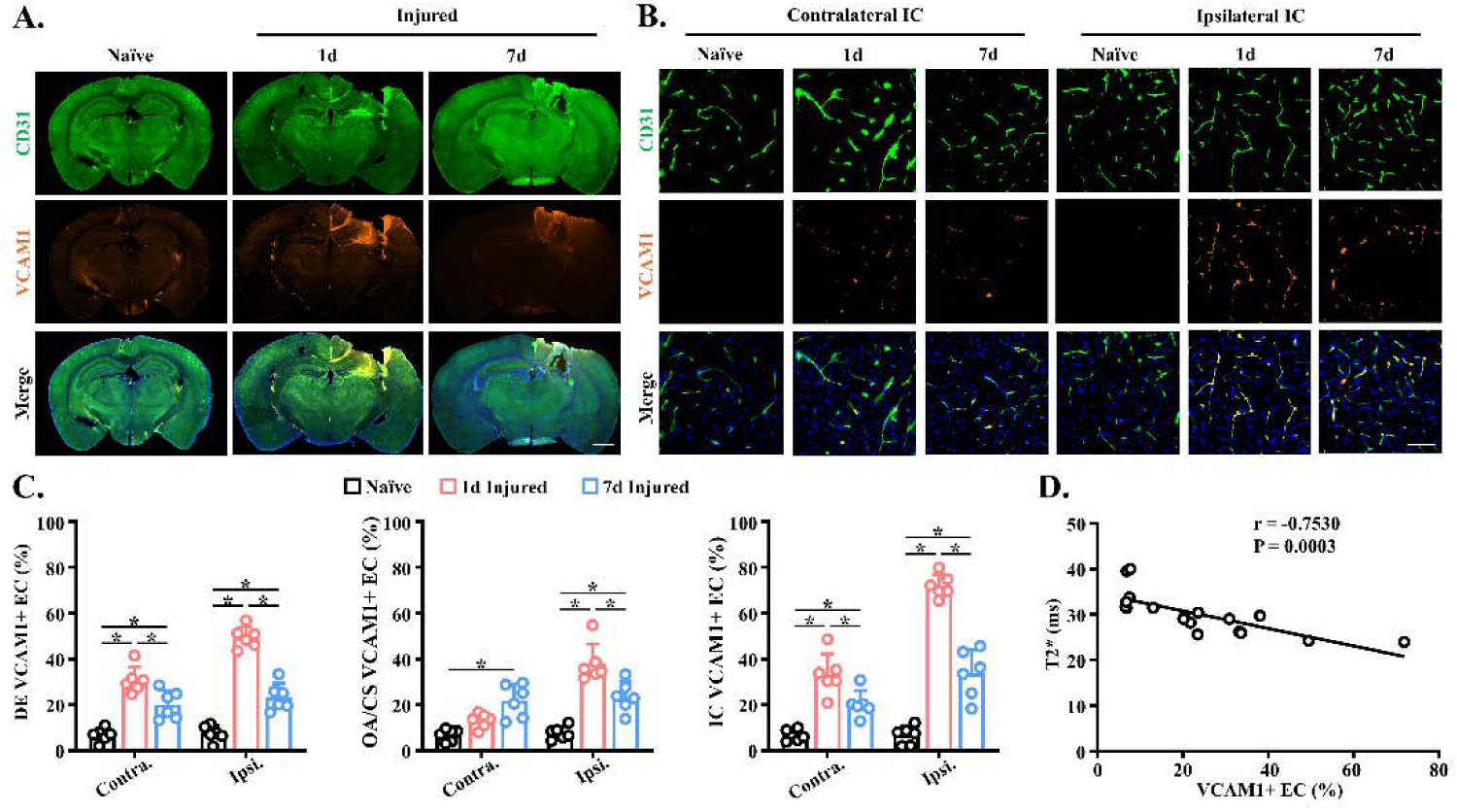
VCAM1 expression correlates with MoNP-SPION MRI contrast across time and brain regions after TBI. (A) Representative whole brain IF images of CD31, VCAM1, and nuclei from naïve, 1-day injured, and 7-day injured mice; scale bar = 1 mm. (B) Representative magnified images of CD31 and VCAM1 colocalization within the IC of the contralateral and ipsilateral hemisphere in naïve, 1-day injured, and 7-day injured mice; scale bar = 50 µm. (C) Quantification of VCAM1-positive ECs within the DE (left), OA/CS (center), and IC (right) of the contralateral and ipsilateral hemispheres in naïve, 1-day injured, and 7-day injured mice; n = 6. (D) Scatter plot and linear regression of T2* values against their spatiotemporally corresponding VCAM1 expression values. All data is presented as mean ± SD. * indicates p < 0.05. Statistical significance was calculated with two-way ANOVA with Tukey’s post-hoc multiple comparisons, Pearson’s correlation analysis was performed to assess relationships between variables.

### 2.6 MoNP-SPION detects therapeutic modulation of cerebrovascular inflammation post-TBI

To determine whether MoNP-SPION can monitor therapeutic response, we tested two agents, Mc and At, both of which have demonstrated pleiotropic anti-inflammatory effects *in vitro* and in preclinical TBI models (**Fig. 6A**) (*23*, *24*). Western blot analysis showed that both agents suppressed TNFα-induced VCAM1 expression in HBMECs (**Fig. S11**). To assess whether these effects translate *in vivo*, CCI-injured mice received daily intraperitoneal injections of Mc or At for 7 days. IF analysis of brain sections confirmed that Mc markedly reduced VCAM1 expression, while At produced modest reductions relative to vehicle controls (**Figs. 6B & 6C**). Consistent with these findings, *in vivo* MoNP-SPION-enhanced MR imaging at day 7 showed persistent hypointense contrast throughout the brains of vehicle-treated mice, whereas At treatment resulted in mild contrast reduction and Mc treatment largely eliminated hypointense contrast (**Fig. 6D**). Correspondingly, quantitative T2* decay analysis showed more rapid signal recovery in Mc-treated mice compared to the vehicle- and At-treated groups (**Fig. 6E**, **Table 3**). *Ex vivo* imaging further supported these findings, revealing substantially reduced MoNP-SPION-associated contrast in Mc-treated brains (**Fig. 6F**). Regional T2* quantification demonstrated only marginal recovery following At treatment, whereas Mc induced significant T2* recovery across most brain regions compared to vehicle controls (**Fig. 6G, Table S3**). Collectively, these results demonstrate that MoNP-SPION-enhanced MR imaging sensitively detects therapeutic suppression of cerebrovascular inflammation following TBI.

**Figure 6.**
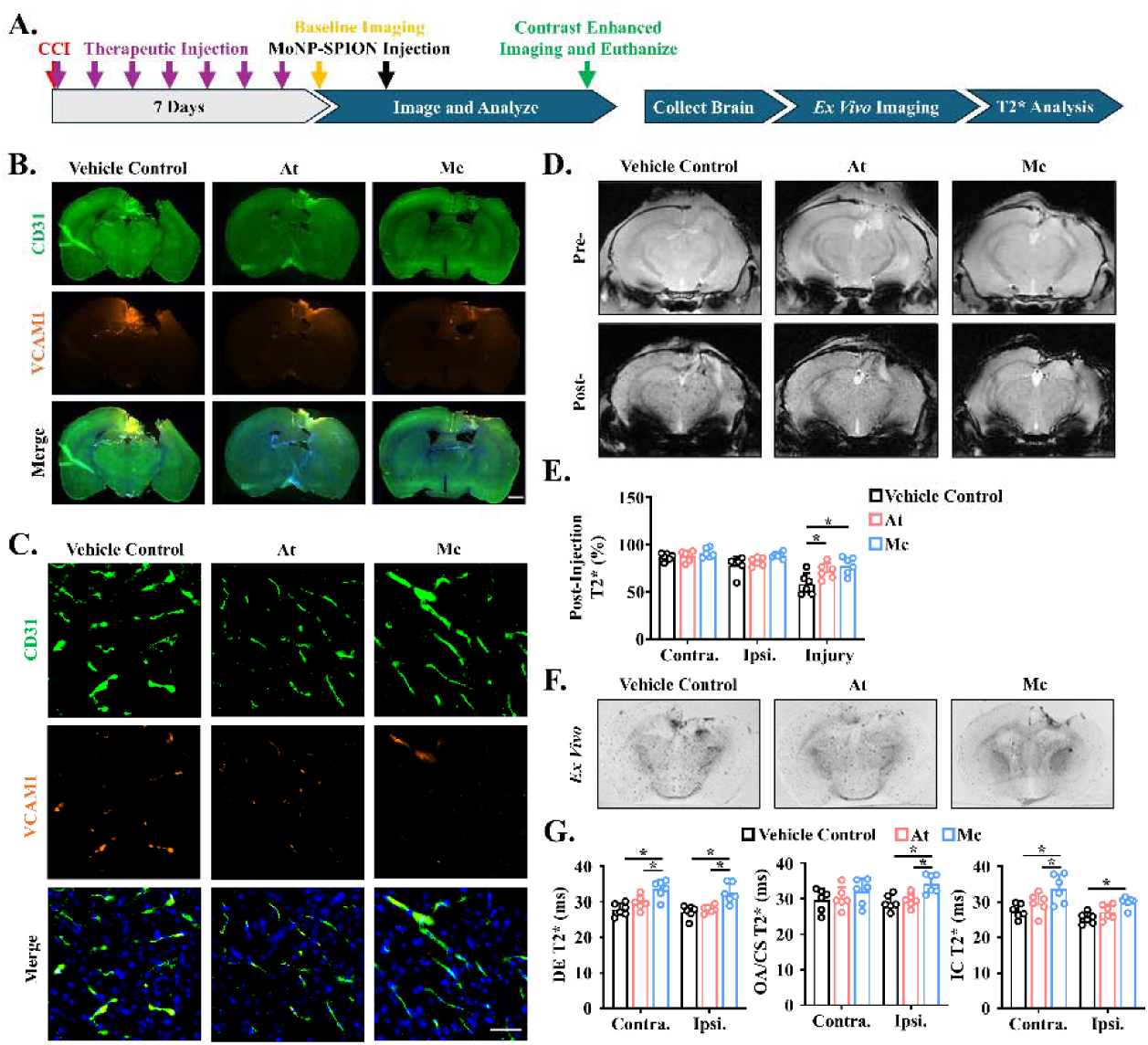
Anti-inflammatory treatment suppresses post-TBI vascular inflammation and associated MRI contrast. (A) Schematic diagram of therapeutic treatment and MR imaging timeline. (B) Representative whole brain IF images of CD31, VCAM1, and nuclei from vehicle-, Atorvastatin (At)-, and Minocycline (Mc)-treated mice 7 days after CCI; scale bar = 1 mm. (C) Representative magnified immunofluorescent images of CD31 and VCAM1 colocalization within the ipsilateral IC; scale bar = 50 µm. (D) Representative pre- and post-injection T2*-weighted images of injured mouse brains after 7 days of Mc, At, or vehicle control treatment. (E) T2* quantification of the contralateral hemisphere, ipsilateral hemisphere, and injury penumbra as a percentage of baseline values; n = 6. (F) Representative T2*-weighted *ex vivo* MR images of injured mouse brains after 7 days of Mc, At, or vehicle control treatment. (G) T2* quantification of the DE (left), OA/CS (center), and IC (right) within the contralateral and ipsilateral hemispheres of injured mice after 7 days of Mc, At, or vehicle control treatment; n = 6. All data is presented as mean ± SD. * indicates p < 0.05. Statistical significance was calculated with two-way ANOVA with Tukey’s post-hoc multiple comparisons test.

**Table 3.** *In vivo* T2* analysis after 7-day anti-inflammatory treatment. Post-injection T2* decay constants represented as a percentage of pre-injection values across the contralateral hemisphere, ipsilateral hemisphere, and injury penumbra from mouse brains after administration of MoNP-SPION in injured mice after seven days of administration of Mc, At, or a vehicle control; n = 6. Data is represented as mean ± SD.

| Post-Injection<br>T2* (%) |  | Contralateral | Ipsilateral | Injury 869<br>Penumbra |
| --- | --- | --- | --- | --- |
|  | Vehicle<br>Control | 86.83±3.84 | 78.03±9.34 | 58.39±11.14 |
|  | At | 87.87±5.44 | 81.75±4.36 | 71.86±8.24 |
|  | Mc | 91.07±4.99 | 88.62±3.13 | 77.81±9.26 |

## 3. DISCUSSION

Common MRI imaging approaches, including DCE-MRI and DWI, detect overt clinical outcomes such as intensified BBB permeability and white matter lesion development, but fall short of accurately capturing dynamic changes in cerebrovascular inflammation (*11*, *12*). Here, we establish MoNP-SPION as a biomimetic nanoprobe capable of providing quantitative, spatiotemporally resolved molecular imaging of cerebrovascular inflammation following TBI. With respect to targeting specificity and quantitative binding, MoNP binding to inflamed brain ECs scaled with their activation state, enabling accumulation not only within injury penumbra but also distal vascular regions through integrin-β1-VCAM1 interactions (**Fig. 1**). Incorporating SPION translated this targeting behavior into a quantitative MRI readout, generating hypointense contrast proportional to the degree of endothelial activation and nanoparticle uptake, while also providing a rapid imaging window and subsequent tissue clearance (**Fig. 2**). Compared to non-targeted counterpart, MoNP-SPION produced spatially heterogeneous reductions in T2* decay across the injury plane as well as rostral and caudal regions (**Fig. 3**). This effect was found to be partially recovered, but not completely, after seven days, demonstrating sensitivity to both spatial and temporal heterogeneity of post-TBI inflammation (**Fig. 4**). The specificity of MoNP-SPION contrast as a readout of cerebrovascular inflammation was confirmed by a strong correlation between regional VCAM1 expression and T2* decay constants after TBI (**Fig. 5**). Its sensitivity was further demonstrated by the ability to detect differential attenuation of cerebrovascular inflammation following therapeutic intervention, with Mc producing marked contrast recovery and At showing modest effects (**Fig. 6**). Collectively, this MoNP-SPION-enhanced MRI approach addresses key limitations of standard imaging methods by enabling efficient and selective endothelial targeting with molecularly grounded, dynamic readouts of cerebrovascular inflammation.

Our previous work established MoNP-SPION as an effective biomimetic contrast agent for imaging and assessing atherosclerotic formation, where monocyte membrane cloaking conferred multivalent targeting of inflamed endothelium lining the plaque surface (*18*). Extending this platform to the cerebrovascular compartment addresses a clinical unmet need as existing imaging tools lack the sensitivity to directly capture molecular alterations of the endothelium underlying cerebrovascular inflammation in TBI. Endothelial activation and subsequent adhesion molecule expression are well recognized as a continuum processes rather than binary “on/off” events, with expression levels scaling according to the magnitude and duration of inflammatory stimuli (*25*, *26*). Our *in vitro* findings are consistent with this paradigm; we observed that both MoNP and MoNP-SPION binding to brain ECs increased in proportion to TNFα-induced activation, until reaching maximal VCAM1 density. Importantly, this saturation-dependent profile suggests that *in vivo*, regional differences in MoNP accumulation reflect genuine differences in endothelial activation magnitude, lending biological meaning to the spatial heterogeneity observed in subsequent MRI experiments. *In vivo*, this enabled the dissemination of MoNP to more distal regions of the brain whereas uncoated NP remained confined to the immediate injury site, potentially through passive leakage through damaged and ruptured vasculature rather than an active targeting effect (*27*). These findings indicate that TBI-induced cerebrovascular inflammation extends beyond the lesion core and may be underestimated by structural imaging alone. This specificity is consistent with the well-established VLA4-VCAM1 monocyte adhesion axis (*21*); blocking β1 integrin, which forms VLA4 on the MoNP surface, abolished distal accumulation and confined signal to the injury penumbra, confirming that MoNP targeting is receptor-mediated and not merely a consequence of passive barrier disruption.

Conventional iron oxide formulations, including ferumoxytol, require delayed imaging windows of up to 24 hours to achieve sufficient contrast, complicating longitudinal workflows and limiting clinical practicality (*15*). This limitation largely reflects their dependence on passive accumulation, an inherently inefficient and nonspecific process. In contrast, by actively targeting inflamed endothelium, MoNP-SPION circumvents this constraint and achieves peak contrast as early as 3 hours post-injection, enabling a substantially early and more practical imaging workflow. Beyond imaging efficiency, signal recovery toward baseline after 9 hours further suggests relatively rapid clearance and facilitating longitudinal re-imaging. This relatively short brain retention window may also be advantageous in limiting prolonged local iron exposure, reducing the risk of iron-mediated oxidative stress and pathological protein aggregation (*3*, *28*). It is worth nothing that all quantitative MRI readouts in this study are based on voxel-wise T2* decay constants rather than single-echo signal intensity measurements. Many SPION-enhanced MRI studies rely predominantly on visual intensity-based endpoints, which are susceptible to saturation effects and constrained dynamic range, limiting their ability to resolve graded differences in probe burden (*29*, *30*). Conversely, T2* decay reflects local magnetic susceptibility and scales with iron oxide concentration, providing a more quantitative, concentration-dependent readout that strengthens the biological interpretation of spatial contrast heterogeneity across all subsequent experiments (*31*).

Utilizing this quantitative T2* framework, we found that MoNP-SPION induced a spatially heterogenous contrast enhancement extending well beyond the primary injury site, with the strongest T2* decay observed within the ipsilateral hemisphere. One day post-TBI, a critical window for secondary injury progression, MoNP-SPION produced a robust, speckled hypointense signal throughout a 4 mm longitudinal span across both hemispheres. While the greatest signal reductions were observed at the ipsilateral epicenter, significant contrast was also detected in rostral and caudal regions, revealing a diminishing yet persistent inflammatory gradient. In comparison, uncoated P-SPION produced minimal contrast outside of the injury penumbra. These observations are consistent with our fluorescent MoNP findings and with prior report showing that BBB disruption and passive leakage, measured by K^trans^, are often restricted to the superficial cortical layers near the impact site (*32*). The persistence of the granular, speckled contrast pattern further supports the mechanism of active targeting over parenchymal diffusion. Our findings suggest that compared to structural imaging and DCE-MRI, MoNP-SPION-enhanced MRI map a much broader “molecular penumbra” after TBI. By mimicking the innate vascular targeting behavior of monocytes as inflammatory signaling propagates through the cerebrovascular tree, this platform provides a functional readout of regions that appear structurally intact but are biochemically “primed” by inflammatory signaling.

Importantly, the neuroinflammatory response to TBI is a dynamic and regionally evolving process that unfolds over an extended period (*2*, *7*). Consistent with this well-characterized pathophysiology, our longitudinal, spatially mapped MRI analyses showed an overall reduction in MoNP-SPION contrast at 7 days post-injury compared with 1 day, supporting partial resolution of cerebrovascular inflammation (*33–35*). Regional *ex vivo* high resolution T2* analysis across a coronal section of the brain (DE, OA/CS, and IC) further indicated heterogeneous signal rather than uniform. Notably, the contralateral OA/CS maintained significantly reduced T2* values relative to naïve controls at 7 days, indicating persistent MoNP-SPION-associated contrast in this distal region. Histological assessment of VCAM1 expression independently supported this imaging finding, as the contralateral OA/CS was the only region that maintained elevated inflammatory marker expression while other regions showed clear reduction by day 7 compared to 1 day post-injury. This persistent distal signal suggests that post-TBI inflammation may continue to propagate beyond the lesion core, potentially driven by sustained cytokine and chemokine signaling or secondary vascular activation through vulnerable distal microvascular bed (*2*, *6*, *7*). The strong linear correlation between the spatiotemporal presence of VCAM1-positive ECs and quantitative T2* decay readouts (i.e., r = -0.7530) further establishes MoNP-SPION contrast as a high-fidelity readout of cerebrovascular inflammation. By detecting these inflammatory signatures at day 7, our MoNP-SPION platform reveals distal, localized pockets of persistent cerebrovascular stress that would likely escape conventional imaging, thereby providing a more complete inflammatory map of the evolving cerebrovascular injury after TBI.

Beyond tracking longitudinal and regional dynamics of cerebrovascular inflammation post-TBI, a clinically useful molecular imaging probe must be sensitive enough to detect treatment-induced changes. To this end, we evaluated two clinically relevant therapeutic agents, Mc and At, both of which have been found to possess potent anti-inflammatory properties and have previously shown efficacy in mitigating post-traumatic cerebrovascular dysfunction (*23*, *24*). Although our *in vitro* assessment confirmed that both agents suppressed the cytokine-induced endothelial activation, MoNP-SPION-enhanced MRI of post-TBI mice revealed clearly divergent *in vivo* therapeutic responses. Following a 7-day treatment regimen, Mc produced a marked reduction in MoNP-SPION accumulation throughout the entire brain, reflected by lower T2* decay values across all regions of the cortex relative to vehicle controls. These findings indicate that Mc treatment effectively diminished the TBI-triggered cerebrovascular endothelial activation. Conversely, At administration resulted in only modest attenuation of MoNP-SPION-induced contrast, with limited recovery in T2* values compared to vehicle controls. Despite shared anti-inflammatory effects *in vitro*, this divergence between Mc and At is consistent with prior clinical observations that At often requires longer-term administration before meaningful reductions in inflammatory biomarkers become apparent (*36*). In addition, our results align with prior molecular imaging studies targeting VCAM1 expression in stroke models, in which At produced only partial recovery of T2* contrast (*30*). The ability of MoNP-SPION-enhanced MRI to distinguish the robust anti-inflammatory effect of Mc from the more limited short-term response to At highlights the sensitivity of this platform to pharmacologic differences in treatment response. As such, the MoNP-SPION platform may serve as a noninvasive approach for monitoring therapeutic efficacy and refining treatment timing for TBI.

While our current study provides important insight into the dynamic imaging capacity of the MoNP-SPION platform in the cerebrovascular bed, several limitations must be addressed before clinical translation can be fully considered. Notably, the CCI was conducted at a single injury severity, but whether MoNP-SPION-enhanced MRI can sensitively detect mild TBI phenotypes remains unknown; it will be particularly important to define the threshold of detection sensitivity *in vivo* to assess the full range of translational utility (*37*). In this study, VCAM1 was used as the sole representative marker to assess cerebrovascular inflammation but this alone cannot capture the full molecular and cellular complexity of the post-TBI vascular responses. In addition, endothelial inflammatory programs evolve over time and regulate the staged recruitment of different leukocyte populations, such as neutrophils and T-lymphocytes within distinct time frames (*2*). Future studies considering membranes derived from these alternative immune cell populations may help tailor the imaging platform toward specific inflammatory phases or diagnostic windows. Although we showed that fluorescent MoNP colocalized with inflamed cerebral ECs in the TBI-brain, it remains unknown whether MoNP-SPION undergoes transcytosis following EC uptake, and to what extent SPION deposition occurs within the brain parenchyma beyond the cerebrovascular compartment. Lastly, given well-established sex differences in TBI neuroinflammation and recovery trajectories, future work should determine whether MoNP-SPION targeting and signal profiles differ between sexes (*38*).

## 4. CONCLUSION

In summary, we establish MoNP-SPION as the first clinically translatable biomimetic nanoprobe for quantitative imaging of spatially and temporally heterogeneous cerebrovascular inflammation after TBI. Through β1-mediated interactions with endothelial surface receptors, MoNP binding scales with endothelial activation, providing a molecularly specific readout of cerebrovascular inflammation. By integrating monocyte-mimicking vascular targeting with SPION-based MRI contrast, this platform enables longitudinal assessment of TBI-induced vascular dysfunction and sensitive detection of vascular recovery following therapeutic intervention. In doing so, MoNP-SPION reveals the broader inflammatory footprint of TBI, uncovering activated cerebrovascular territories that would otherwise remain undetected by conventional imaging approaches. Together, these features position MoNP-SPION as a promising translational platform for real-time, noninvasive monitoring of cerebrovascular inflammation in TBI and other neurological disorders involving vascular dysfunction, including stroke and vascular dementia.

## 5. MATERIALS AND METHODS

### 5.1 Controlled Cortical Impact (CCI) Model

All animal experiments performed in this study have been approved by the Institutional Animal Care and Use Committee (IACUC) of Arizona State University (23-1998R and 26-2173R). Male and female C57BL/6 mice (8-10 weeks old; Jackson Laboratory) were subjected to a well-established CCI model to induce a moderate unilateral TBI over the right somatosensory cortex (*39*). Briefly, mice were anesthetized with isoflurane (3% induction, 1.5% maintenance) and secured on a stereotaxic frame. A 10 mm midline incision exposed the skull and a 3 mm biopsy punch was used to perform a craniotomy centered at 1.5 mm posterior to bregma and 1.5 mm lateral to midline. The CCI was induced at 6.0 m/s for a duration of 100 ms at a depth of 1.5 mm using a Leica Impact One (Leica Biosystems) device with a 2 mm probe. After the bleeding was controlled, the skin was sutured and antibiotic ointment was applied to the suture site. Subcutaneous injections of buprenorphine (0.5 mg/kg) and sterile saline (0.5 mL) were administered, and mice were taken off anesthesia. Mice were placed in a fresh cage on a warming pad and monitored for 1 hour prior to being transferred to the vivarium.

### 5.2 Cell Culture

HBMECs (Cell Systems, ACBRI376) was maintained in complete classic medium under standard cell culture conditions (37 °C, 5% CO2, 100% humidity). For monocytes, bone marrow cells were isolated and maintained in RPMI 1640 medium supplemented with 20 ng/ml of MCSF under standard cell culture conditions for 5 days. Monocytes were then magnetically sorted and their plasma membranes were isolated using Minute Plasma Membrane Isolation kit (Invent Biotechnologies).

### 5.3 Western Blot

To assess heterogeneity in cerebrovascular EC activation, HBMECs were incubated with increasing concentrations of TNFα (0-25 ng/ml) for 3 hours. To assess therapeutic effects on HBMEC activation, cells were pretreated overnight with either 50 µM Mc (G-Biosciences, #GBM-2568), 100 µM At (Cayman Chemical, #C833M82), 2% dimethyl sulfoxide (DMSO), or saline for 24 hours prior to treatment with TNFα at a concentration of 10 ng/ml. After TNFα activation of all cells, they were lysed and proteins were collected with RIPA buffer. Western blot was then performed to probe for VCAM1 (Cell Signaling, #13622, 1:1000) and GAPDH (Cell Signaling, #2118, 1:1000) and imaged with a gel documentation system (Analytik Jena).

### 5.4 Nanoparticle Formulation and Characterization

MoNP and MoNP-SPION were synthesized as previously described (*18*, *20*). Briefly, to generate fluorescent MoNP, PLGA (Sigma-Aldrich) and lipophilic carbocyanine dye DiD (Biotium) were dissolved in dichloromethane (DCM) (Sigma-Aldrich) and emulsified in 2% poly(vinyl alcohol) (PVA) before allowed to dry for 3 hours in a 0.5% PVA solution to form nanoparticle cores (NP). For MoNP-SPION, 1 mg SPION (Sigma-Aldrich) resuspended in ultrapure water was emulsified with PLGA dissolved in DCM and then further emulsified within 0.5% PVA to generate SPION-loaded polymeric cores (P-SPION). Both NP and P-SPION were washed with pure water at 17,000g. Plasma membrane fractions were coated onto the surface of NP and P-SPION at a 1:10 (w/w) ratio using an ultrasonic bath. DLS (Zetasizer, Malvern Analytical) was used to measure physicochemical properties of each nanoparticle. To assess morphology and SPION encapsulation of MoNP-SPION, negative staining using 0.5% uranyl acetate was conducted to visualize with TEM (Talos L120C, Thermo Fisher Scientific) at the ASU Eyring Materials Center.

### 5.5 *In Vitro* Assessment of Nanoparticle Binding and Uptake

HBMECs were treated with 10 ng/ml of TNFα for 3 hours to induce inflammatory activation. Activated HBMECs were then incubated with fluorescently labeled NP or MoNP at 50 µg/ml, or with P-SPION or MoNP-SPION at 10 µg Fe/ml, for 30 minutes. Cells treated with fluorescent MoNP were then fixed with 4% paraformaldehyde (PFA) before being imaged on a confocal microscope (Leica Stellaris 8) to confirm intracellular uptake. HBMECs treated with MoNP-SPION were fixed and stained with 5% potassium ferrocyanide and 10% HCl and imaged using a Lionheart FX automated microscope (Agilent) to visualize iron deposition. To assess nanoparticle interaction under varying inflammatory states, HBMECs were treated with increasing concentrations of TNFα and then incubated with fluorescently labeled MoNP or MoNP-SPION for 10 minutes. Cells treated with MoNP were then either processed for fluorescence microscopy or flow cytometry. For microscopy, cells were fixed with 4% PFA and imaged to measure total fluorescent intensity. For flow cytometry, cells were collected, fixed, and measured for DiD fluorescence (Attune NxT, ThermoFisher). For cells treated with MoNP-SPION, iron was directly quantified in each cell culture by first digesting in 20% HCl overnight. To the HCl solution, 5% potassium ferrocyanide was added at a 1:1 (v/v) ratio and the absorbance was measured using a plate reader (Synergy H1, BioTek) at 650 nm. MoNP-SPION was further measured through phantom imaging wherein cells were collected in PBS and suspended within a 1% agarose gel. T2*-weighted MR imaging (TE: 16.9; TR: 833; α: 60) was conducted using a preclinical 9.4T MRI system (MRSolutions) to assess contrast changes in each phantom.

### 5.6 IVIS Imaging

RO injections were used to administer saline or a 10 mg/kg dose of NP or MoNP 24 hours post-injury. Three hours post-injection, mice were euthanized via intraperitoneal (IP) injection of a lethal dose of sodium pentobarbital (200 mg/kg Euthasol) and transcardially perfused with PBS and 4% PFA. The brain was collected into 4% PFA overnight for fixation prior to imaging using IVIS (PerkinElmer). For VLA4 blocking, MoNP was incubated with anti-β1 (Cell Signaling, #4706, 1:50) or IgG (Santa Cruz, SC-2025, 1:50) for 30 minutes prior to mice receiving an RO dose at 24 hours post-CCI. Three hours after injection, mice were euthanized, and tissue was collected and fixed for IVIS imaging. Spectral unmixing was used to isolate DiD fluorescent signal from tissue autofluorescence. Regions of interest (ROIs) were placed over the entire brain to measure fluorescent signal in the entire tissue.

### 5.7 Animal Magnetic Resonance Imaging

For all *in vivo* MR imaging experiments, mice were anesthetized and secured in the mouse head coil. While secured in the 9.4T MRI, mice were anesthetized with isoflurane (3% induction, 1.5% maintenance) and warmed with continuous warm air flow. For serial imaging (TE: 4.8-61.5; TR: 833; α: 45; Slice Thickness: 1 mm), after baseline images were acquired, mice were administered RO doses of MoNP-SPION at 5 mg Fe/kg, 24 hours post-injury. Subsequent imaging was conducted at 1, 3, 6, 9, and 24 hours post-injection. After the completion of the 24-hour imaging timepoint, mice were euthanized. Comparative imaging (TE: 4.8-61.5; TR: 600; α: 45; Slice Thickness: 0.5 mm) was then conducted by collecting baseline imagines, followed by RO administration of P-SPION or MoNP-SPION at 5 mg Fe/kg, or saline, at 24 hours post-injury. Subsequent images were acquired at 3 hours post-injection. Slice thickness of 0.5 mm was determined to assess more discrete spatial discrepancies within the injury penumbra than what was acquired at 1 mm. After imaging was completed, mice were euthanized and major organs were collected. Brains were perfused and fixed prior to embedding within 1% agarose for *ex vivo* phantom MR imaging. For longitudinal assessment and therapeutic recovery studies, after sustaining TBI, mice were randomly assigned to groups to be administered intraperitoneal doses of either Mc (45 mg/kg), At (1 mg/kg), DMSO (2%) as a vehicle control, or saline every 24 hours for 7 days. Seven days post-injury, baseline MR images were acquired (TE: 4.8-61.5; TR: 600; α: 45; Slice Thickness: 0.5mm). After baseline images were acquired, mice were administered a 5 mg Fe/kg RO dose of MoNP-SPION and reimaged after 3 hours. Once imaging was completed, mice were euthanized and their major organs were collected. Brains were collected for *ex vivo* MR imaging. Mice administered saline during the longitudinal study were used for temporal comparisons to naive and 1-day injured mice. The same group was then used to assess 7-day post-injury VCAM1 expression patterns. All MR images were acquired using a T2*-weighted multigradient echo sequence.

### 5.8 Histology

Organs were fixed overnight in 4% PFA and subsequently transferred through a sucrose gradient over several days. Tissues were embedded in OCT compound, flash-frozen, and sectioned using a cryostat (Leica). Brain sections were collected from the lesion core. For IF analysis, sections were permeabilized with 0.1% Triton X-100, washed with PBS, and stained for VCAM1 (Thermo, #MA5-11447, 1:100) and CD31 (BD Bioscience, #553370, 1:100). For colorimetric staining of iron, sections were incubated in a 1:1 solution of 5% potassium ferrocyanide and 20% HCl for 30 minutes, followed by PBS washes and counterstaining with nuclear fast red (Sigma) for 1 minute. After staining, slides were dehydrated through a graded ethanol series followed by xylene, then mounted using standard mounting media. All histological imaging was performed using a slide scanner (VS200, Olympus).

### 5.9 Image Analysis

The cerebral cortex was segmented into anatomically defined regions based on proximity to the lesion core. Dorsal analysis of the cerebral cortex was restricted to the IC, while ventral regions included the OA/CS. The DE, containing the thalamus and hypothalamus, was similarly analyzed. Each anatomical region was further subdivided by hemisphere, resulting in six distinct analysis regions: contralateral DE, contralateral OA/CS, contralateral IC, ipsilateral DE, ipsilateral OA/CS, and the ipsilateral IC. Region boundaries were guided by the Allen mouse brain atlas. To avoid signal saturation of the cortex at the injury, all measurements were acquired outside of the immediate injury penumbra. IF stained brain sections located within the injury plane were analyzed using ImageJ. Image thresholds were applied to delineate CD31-positive and VCAM1-positive vasculature from the surrounding tissue autofluorescence, after which VCAM1-positive area was then measured as a percentage of total CD31-positive area (*40*).

To calculate T2* decay constants from MR images, SI across all TEs were first fitted on a voxel-wise basis to the following mono-exponential model (*41*):

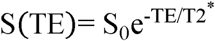

where S(TE) is the SI at a given TE, S_0_ is the initial SI of the first TE, and T2* is the effective transverse magnetization decay constant. T2* maps were then derived from the exponential fit with voxel-specific decay constants calculated as the time at 1/e (37%) signal, and R2* maps were calculated as the reciprocal (R2* = 1/T2*) (*42*). ROIs were manually defined and applied consistently across all TEs. For all *in vivo* MR images, T2* decay was only measured within each hemisphere and the cerebral cortex of the injury penumbra due to artifacts from motion and susceptibility-induced signal voids originating in the ear canal. To quantify the *in vivo* changes associated with nanoparticle accumulation, pre- and post-injection T2* decay constants were compared and measured as a percent change:

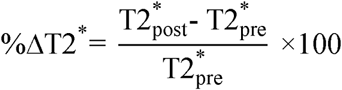

where 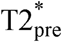 and 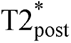 are T2* decay constant before or after nanoparticle administration respectively. For *ex vivo* MR imaging, ROIs were selected within the contralateral IC, contralateral OA/CS, contralateral DE, ipsilateral IC, the ipsilateral OA/CS, and ipsilateral DE regions of brain slices within the injury plane for direct quantification of regional T2* decay constants.

### 5.10 Statistical Analysis

All graphical data is presented as mean ± standard deviation (SD). Statistical analysis from at least three biological replicates was performed using GraphPad Prism 2025. One- and two-way ANOVA were performed for the comparison of multiple groups followed by Tukey’s multiple comparisons post-hoc test. For comparison between two groups, a two tailed unpaired t-test was used. For correlation analysis of VCAM1-positive EC and T2* decay, datapoints from six individual biological replicates in all six brain regions (ipsilateral DE, ipsilateral OA/CS, ipsilateral IC, contralateral DE, contralateral OA/CS, and contralateral IC) at each timepoint (naïve, 1d, 7d) were averaged and plotted against each other, generating 18 distinct datapoints. Correlation between T2* and VCAM1 expression was conducted using a linear regression to determine Pearson’s coefficient. A p-value < 0.05 was considered to be statistically significant.

## ACKNOWLEDGEMENTS

The authors acknowledge the use of Regenerative Medicine Core, Advanced Light Microscopy Core, Histology Core, Flow Cytometry Core, and Preclinical Imaging Core facilities at Arizona State University. Schematic diagrams in all figures were created using BioRender.com.

## FUNDING

This work was supported in part by ASU startup funds and NSF CAREER Award 2544114 (to K.-C. W.) and by NIH R01NS116657-05 (to S.E.S.). J.R., A.S., and T.-Y.W. were supported by the Arizona State University Graduate College Completion Fellowship.

## AUTHOR CONTRIBUTIONS

Conceptualization: J.R., A.S., S.E.S., and K.-C.W. Methodology: J.R., A.S., D.O., S.E.S., and K.-C.W. Investigation: J.R., A.S., D.O., M.K., O.A., T.-Y.W., S.E.S., and K.-C.W. Visualization: J.R., A.S., D.O., S.E.S., and K.-C.W. Supervision: S.E.S., and K.-C.W. Writing – Original Draft: J.R. and K.-C.W. Writing – Review & Editing: J.R., A.S., S.E.S., and K.-C.W.

## COMPETING INTERESTS

The authors declare that they have no competing interests.

## DATA AND MATERIALS AVAILABILTY

All data needed to evaluate the conclusions in the paper are present in the paper and/or the Supplementary Materials.

## Notes

### Competing Interest Statement

The authors have declared no competing interest.

## REFERENCES

1. C. A. Taylor, J. M. Bell, M. J. Breiding, L. Xu, Traumatic brain injury-related emergency department visits, hospitalizations, and deaths - United States, 2007 and 2013. MMWR Surveill Summ 66, 1–16 (2017).

2. T. T. Postolache, A. Wadhawan, A. Can, C. A. Lowry, M. Woodbury, H. Makkar, A. J. Hoisington, A. J. Scott, E. Potocki, M. E. Benros, J. W. Stiller, Inflammation in traumatic brain injury. J Alzheimers Dis 74, 1–28 (2020).

3. P. M. A. Muneer, N. Chandra, J. Haorah, Interactions of oxidative stress and neurovascular inflammation in the pathogenesis of traumatic brain injury. Mol Neurobiol 51, 966–979 (2015).

4. D. J. Loane, A. Kumar, Microglia in the TBI brain: the good, the bad, and the dysregulated. Exp Neurol 275 **Pt** **3**, 316–327 (2016).

5. E. A. van Vliet, X. E. Ndode-Ekane, L. J. Lehto, J. A. Gorter, P. Andrade, E. Aronica, O. Gröhn, A. Pitkänen, Long-lasting blood-brain barrier dysfunction and neuroinflammation after traumatic brain injury. Neurobiol Dis 145, 105080 (2020).

6. A. Alam, E. P. Thelin, T. Tajsic, D. Z. Khan, A. Khellaf, R. Patani, A. Helmy, Cellular infiltration in traumatic brain injury. J Neuroinflammation 17, 328 (2020).

7. K. Shi, J. Zhang, J.-F. Dong, F.-D. Shi, Dissemination of brain inflammation in traumatic brain injury. Cell Mol Immunol 16, 523–530 (2019).

8. W. Zhang, D. Xiao, Q. Mao, H. Xia, Role of neuroinflammation in neurodegeneration development. Sig Transduct Target Ther 8, 267 (2023).

9. B. Lee, A. Newberg, Neuroimaging in traumatic brain imaging. NeuroRx 2, 372–383 (2005).

10. C. A. Mutch, J. F. Talbott, A. Gean, Imaging evaluation of acute traumatic brain injury. Neurosurg Clin N Am 27, 409–439 (2016).

11. C. C. Quarles, L. C. Bell, A. M. Stokes, Imaging vascular and hemodynamic features of the brain using dynamic susceptibility contrast and dynamic contrast enhanced MRI. Neuroimage 187, 32–55 (2019).

12. I. D. Croall, V. Lohner, B. Moynihan, U. Khan, A. Hassan, J. T. O’Brien, R. G. Morris, D. J. Tozer, V. C. Cambridge, K. Harkness, D. J. Werring, A. M. Blamire, G. A. Ford, T. R. Barrick, H. S. Markus, Using DTI to assess white matter microstructure in cerebral small vessel disease (SVD) in multicentre studies. Clinical Science 131, 1361–1373 (2017).

13. A.-C. Dupont, B. Largeau, M. J. Santiago Ribeiro, D. Guilloteau, C. Tronel, N. Arlicot, Translocator protein-18 kDa (TSPO) positron emission tomography (PET) imaging and its clinical impact in neurodegenerative diseases. Int J Mol Sci 18, 785 (2017).

14. J. S. Weinstein, C. G. Varallyay, E. Dosa, S. Gahramanov, B. Hamilton, W. D. Rooney, L. L. Muldoon, E. A. Neuwelt, Superparamagnetic iron oxide nanoparticles: diagnostic magnetic resonance imaging and potential therapeutic applications in neurooncology and central nervous system inflammatory pathologies, a review. J Cereb Blood Flow Metab 30, 15–35 (2010).

15. G. B. Toth, C. G. Varallyay, A. Horvath, M. R. Bashir, P. L. Choyke, H. E. Daldrup-Link, E. Dosa, J. P. Finn, S. Gahramanov, M. Harisinghani, I. Macdougall, A. Neuwelt, S. S. Vasanawala, P. Ambady, R. Barajas, J. S. Cetas, J. Ciporen, T. J. DeLoughery, N. D. Doolittle, R. Fu, J. Grinstead, A. R. Guimaraes, B. E. Hamilton, X. Li, H. L. McConnell, L. L. Muldoon, G. Nesbit, J. P. Netto, D. Petterson, W. D. Rooney, D. Schwartz, L. Szidonya, E. A. Neuwelt, Current and potential imaging applications of ferumoxytol for magnetic resonance imaging. Kidney Int 92, 47–66 (2017).

16. S. Martinez De Lizarrondo, C. Jacqmarcq, M. Naveau, M. Navarro-Oviedo, S. Pedron, A. Adam, B. Freis, S. Allouche, D. Goux, S. Razafindrakoto, F. Gazeau, D. Mertz, D. Vivien, T. Bonnard, M. Gauberti, Tracking the immune response by MRI using biodegradable and ultrasensitive microprobes. Sci. Adv. 8, eabm3596 (2022).

17. O. A. Marcos-Contreras, C. F. Greineder, R. Y. Kiseleva, H. Parhiz, L. R. Walsh, V. Zuluaga-Ramirez, J. W. Myerson, E. D. Hood, C. H. Villa, I. Tombacz, N. Pardi, A. Seliga, B. L. Mui, Y. K. Tam, P. M. Glassman, V. V. Shuvaev, J. Nong, J. S. Brenner, M. Khoshnejad, T. Madden, D. Weissmann, Y. Persidsky, V. R. Muzykantov, Selective targeting of nanomedicine to inflamed cerebral vasculature to enhance the blood–brain barrier. Proc. Natl. Acad. Sci. U.S.A. 117, 3405–3414 (2020).

18. J. Rousseau, T.-Y. Wang, S. McClendon, D. Ortega, M. Orlando, S. C. Beeman, B. B. Bartelle, K.-C. Wang, Monocyte-mimetic contrast agent enables targeted and sensitive magnetic resonance imaging of atherosclerotic lesions. Adv Healthc Mater, e02001 (2025).

19. A. Jambusaria, Z. Hong, L. Zhang, S. Srivastava, A. Jana, P. T. Toth, Y. Dai, A. B. Malik, J. Rehman, Endothelial heterogeneity across distinct vascular beds during homeostasis and inflammation. Elife 9, e51413 (2020).

20. H.-C. Huang, T.-Y. Wang, J. Rousseau, M. Orlando, M. Mungaray, C. Michaud, C. Plaisier, Z. B. Chen, K.-C. Wang, Biomimetic nanodrug targets inflammation and suppresses YAP/TAZ to ameliorate atherosclerosis. Biomaterials 306, 122505 (2024).

21. X. Pang, X. He, Z. Qiu, H. Zhang, R. Xie, Z. Liu, Y. Gu, N. Zhao, Q. Xiang, Y. Cui, Targeting integrin pathways: mechanisms and advances in therapy. Sig Transduct Target Ther 8, 1 (2023).

22. A. Jullienne, A. Obenaus, A. Ichkova, C. Savona-Baron, W. J. Pearce, J. Badaut, Chronic cerebrovascular dysfunction after traumatic brain injury. J Neurosci Res 94, 609–622 (2016).

23. P. J. Bergold, R. Furhang, S. Lawless, Treating traumatic brain injury with minocycline. Neurotherapeutics 20, 1546–1564 (2023).

24. H. Wang, J. R. Lynch, P. Song, H.-J. Yang, R. B. Yates, B. Mace, D. S. Warner, J. R. Guyton, D. T. Laskowitz, Simvastatin and atorvastatin improve behavioral outcome, reduce hippocampal degeneration, and improve cerebral blood flow after experimental traumatic brain injury. Exp Neurol 206, 59–69 (2007).

25. J.-J. Chiu, P.-L. Lee, C.-N. Chen, C.-I. Lee, S.-F. Chang, L.-J. Chen, S.-C. Lien, Y.-C. Ko, S. Usami, S. Chien, Shear stress increases ICAM-1 and decreases VCAM-1 and E-selectin expressions induced by tumor necrosis factor-α in endothelial cells. ATVB 24, 73–79 (2004).

26. R. Giri, Y. Shen, M. Stins, S. Du Yan, A. M. Schmidt, D. Stern, K.-S. Kim, B. Zlokovic, V. K. Kalra, β-Amyloid-induced migration of monocytes across human brain endothelial cells involves RAGE and PECAM-1. American Journal of Physiology-Cell Physiology 279, C1772–C1781 (2000).

27. V. N. Bharadwaj, J. Lifshitz, P. D. Adelson, V. D. Kodibagkar, S. E. Stabenfeldt, Temporal assessment of nanoparticle accumulation after experimental brain injury: Effect of particle size. Sci Rep 6, 29988 (2016).

28. Z. Yarjanli, K. Ghaedi, A. Esmaeili, S. Rahgozar, A. Zarrabi, Iron oxide nanoparticles may damage to the neural tissue through iron accumulation, oxidative stress, and protein aggregation. BMC Neurosci 18, 51 (2017).

29. X. Huang, C. Lin, C. Luo, Y. Guo, J. Li, Y. Wang, J. Xu, Y. Zhang, H. Wang, Z. Liu, B. Chen, Fe3O4@M nanoparticles for MRI-targeted detection in the early lesions of atherosclerosis. Nanomedicine: Nanotechnology, Biology and Medicine 33, 102348 (2021).

30. M. Gauberti, A. Montagne, O. A. Marcos-Contreras, A. Le Béhot, E. Maubert, D. Vivien, Ultra-sensitive molecular MRI of Vascular Cell adhesion molecule-1 reveals a dynamic inflammatory penumbra after strokes. Stroke 44, 1988–1996 (2013).

31. F. Bagnato, S. Hametner, E. Boyd, V. Endmayr, Y. Shi, V. Ikonomidou, G. Chen, S. Pawate, H. Lassmann, S. Smith, E. B. Welch, Untangling the R2* contrast in multiple sclerosis: A combined MRI-histology study at 7.0 Tesla. PLoS ONE 13, e0193839 (2018).

32. W. Li, L. Watts, J. Long, W. Zhou, Q. Shen, Z. Jiang, Y. Li, T. Q. Duong, Spatiotemporal changes in blood-brain barrier permeability, cerebral blood flow, T2 and diffusion following mild traumatic brain injury. Brain Res 1646, 53–61 (2016).

33. D. W. Simon, M. J. McGeachy, H. Bayır, R. S. B. Clark, D. J. Loane, P. M. Kochanek, The far-reaching scope of neuroinflammation after traumatic brain injury. Nat Rev Neurol 13, 171–191 (2017).

34. R. Prakash, S. T. Carmichael, Blood-brain barrier breakdown and neovascularization processes after stroke and traumatic brain injury. Curr Opin Neurol 28, 556–564 (2015).

35. M. K. Sköld, C. von Gertten, A.-C. Sandberg-Nordqvist, T. Mathiesen, S. Holmin, VEGF and VEGF receptor expression after experimental brain contusion in rat. J Neurotrauma 22, 353–367 (2005).

36. S. Sabeel, B. Motaung, K. A. Nguyen, M. Ozturk, S. L. Mukasa, K. Wolmarans, D. J. Blom, K. Sliwa, E. Nepolo, G. Günther, R. J. Wilkinson, C. Schacht, A. P. Kengne, F. Thienemann, R. Guler, Impact of statins as immune-modulatory agents on inflammatory markers in adults with chronic diseases: A systematic review and meta-analysis. PLoS One 20, e0323749 (2025).

37. O. N. Kokiko-Cochran, J. P. Godbout, The inflammatory continuum of traumatic brain injury and alzheimer’s disease. Front. Immunol. 9, 672 (2018).

38. A. Simmons, O. Mihalek, H. A. Bimonte Nelson, R. W. Sirianni, S. E. Stabenfeldt, Acute brain injury and nanomedicine: sex as a biological variable. *Front*. Biomater. Sci. 3, 1348165 (2024).

39. V. N. Bharadwaj, C. Copeland, E. Mathew, J. Newbern, T. R. Anderson, J. Lifshitz, V. D. Kodibagkar, S. E. Stabenfeldt, Sex-dependent macromolecule and nanoparticle delivery in experimental brain injury. Tissue Eng Part A 26, 688–701 (2020).

40. H. Yousef, C. J. Czupalla, D. Lee, M. B. Chen, A. N. Burke, K. A. Zera, J. Zandstra, E. Berber, B. Lehallier, V. Mathur, R. V. Nair, L. N. Bonanno, A. C. Yang, T. Peterson, H. Hadeiba, T. Merkel, J. Körbelin, M. Schwaninger, M. S. Buckwalter, S. R. Quake, E. C. Butcher, T. Wyss-Coray, Aged blood impairs hippocampal neural precursor activity and activates microglia via brain endothelial cell VCAM1. Nat Med 25, 988–1000 (2019).

41. D. Milford, N. Rosbach, M. Bendszus, S. Heiland, Mono-exponential fitting in T2-relaxometry: relevance of offset and first echo. PLoS One 10, e0145255 (2015).

42. T. Hesper, H. S. Hosalkar, D. Bittersohl, G. H. Welsch, R. Krauspe, C. Zilkens, B. Bittersohl, T2* mapping for articular cartilage assessment: principles, current applications, and future prospects. Skeletal Radiol 43, 1429–1445 (2014).

