## Supplementary material for "Biomimetic MRI Nanoprobe for Mapping Cerebrovascular Inflammation After Traumatic Brain Injury": Supplimental Data 1

**SUPPLIMENTARY MATERIALS**

**Title**

**Affiliations**

**Field Codes:** Biomedicine and Life Sciences

**SUPPLIMENTARY FIGURES:**

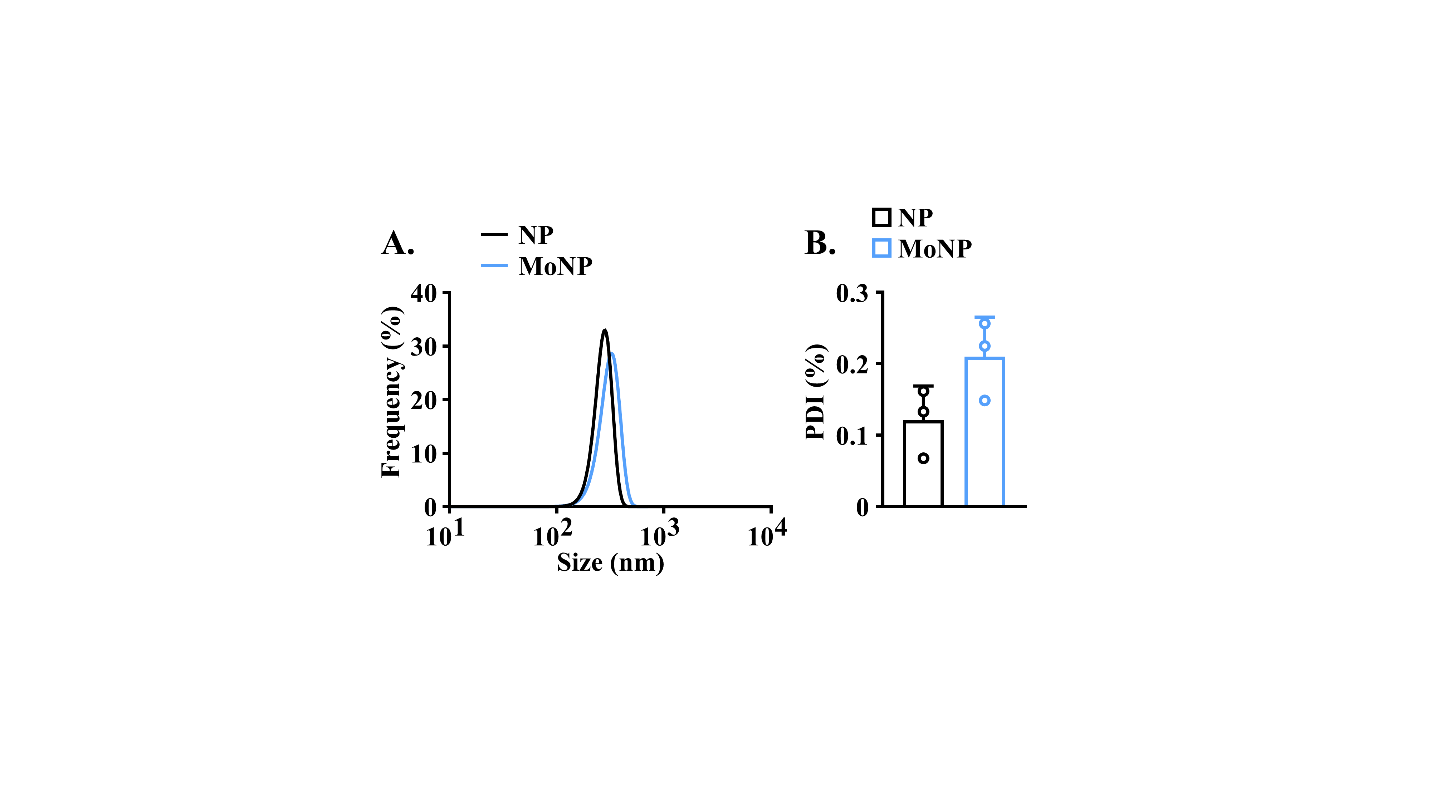

**Figure S1. Physicochemical characterization of fluorescent MoNP.** (A-B) Dynamic light scattering (DLS) measurements of (A) hydrodynamic size and (B) PDI from fluorescent NP and MoNP formulations; n = 3.

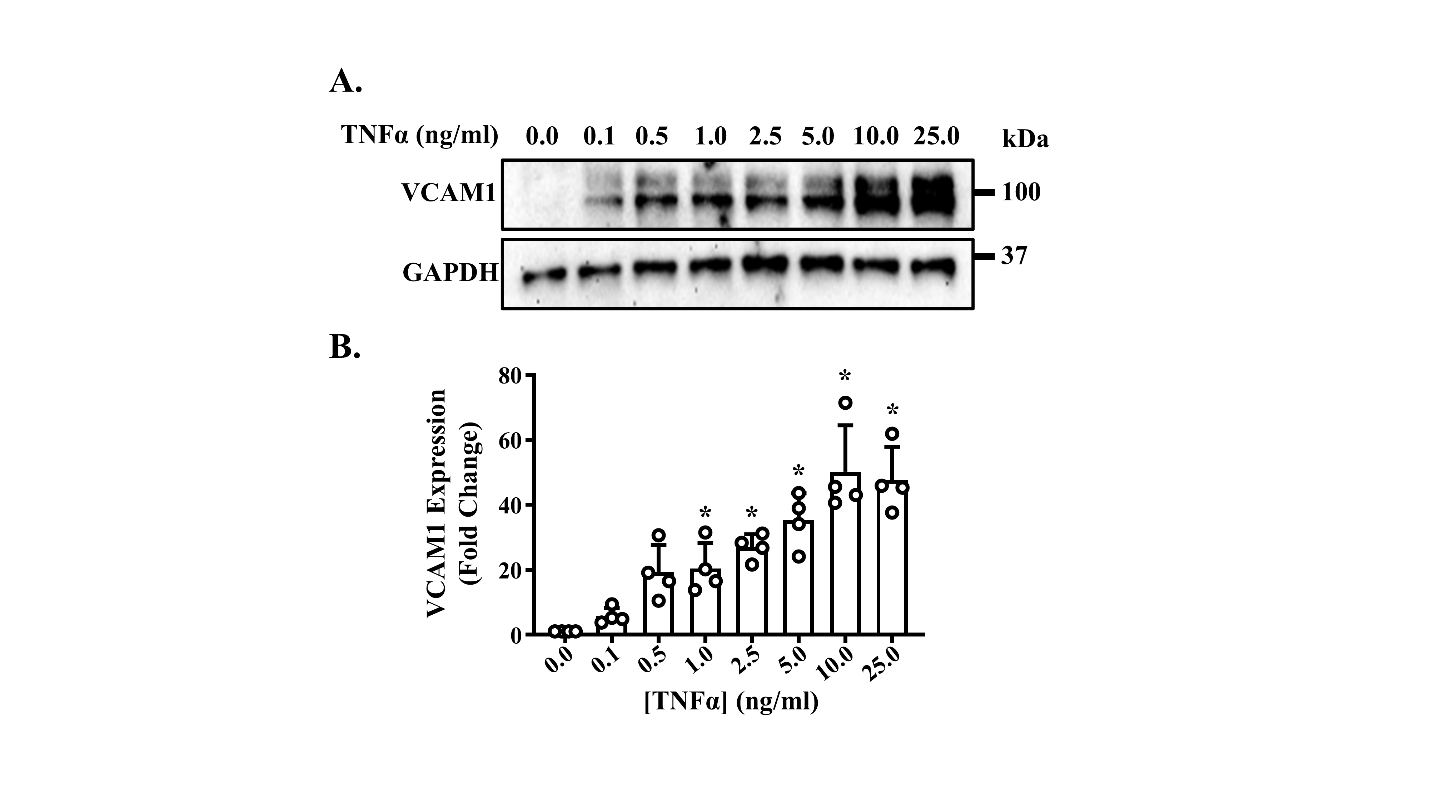

**Figure S2. TNFα-induced VCAM1 expression in HBMECs.** (A) Representative Western blot image of VCAM1 and GAPDH from HBMECs treated with increasing concentrations of TNFα. (B) Quantification of VCAM1 expression; n = 4. Data is presented as mean ± SD. * indicates p < 0.05 compared to untreated control by one-way ANOVA with Tukey's post-hoc multiple comparisons test.

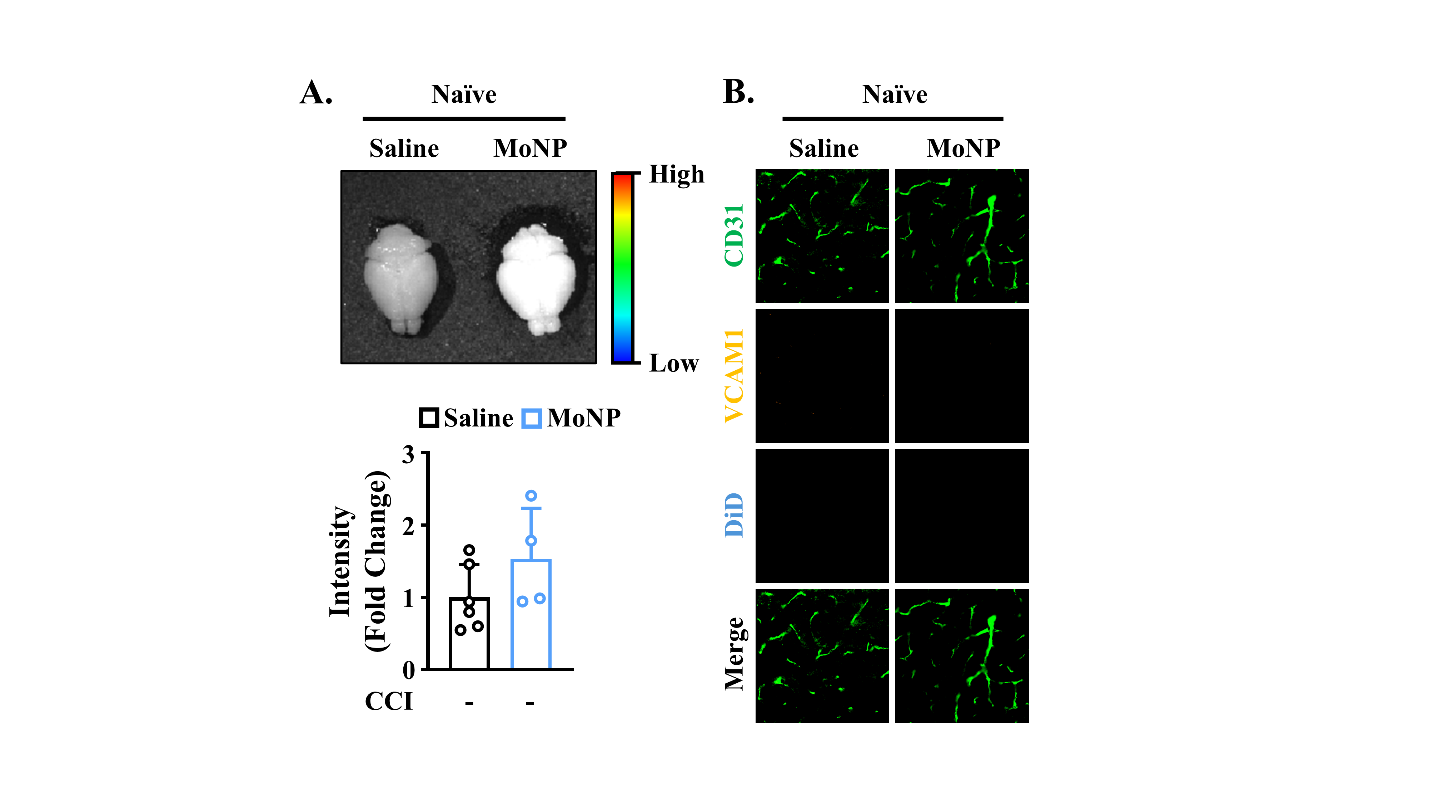

**Figure S3. MoNP accumulation in naïve brains.** (A) Representative IVIS images and quantification of total signal intensity of naïve brains after administration of saline and MoNP; n = 6 for saline and n = 4 for MoNP. (B) Representative IF images of naïve brains after administration of saline and MoNP stained for CD31 (green), VCAM1 (orange), and DiD (cyan). Data is presented as mean ± SD. Statistical significance between two groups was calculated with a two tailed unpaired t-test.

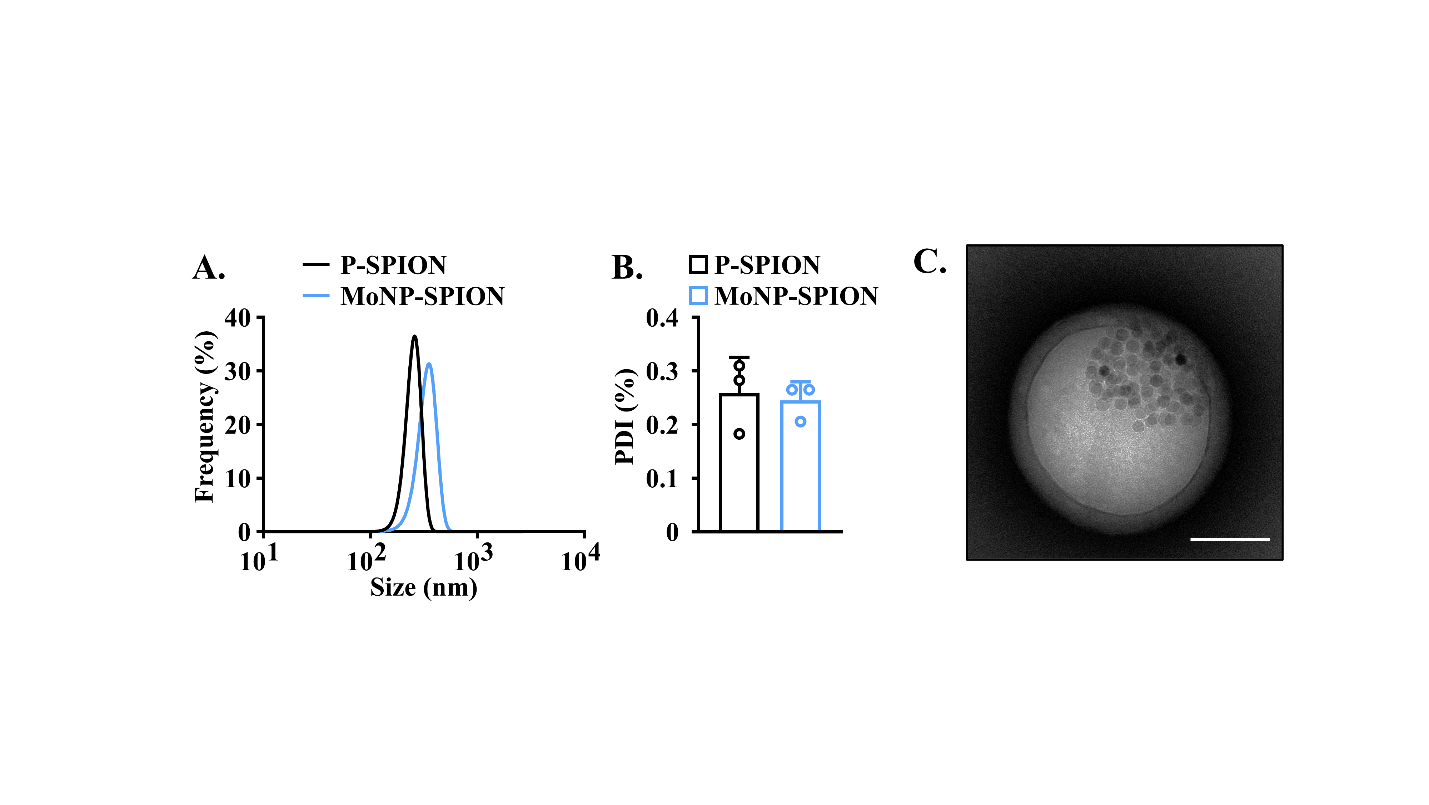

**Figure S4. Physicochemical characterization of MoNP-SPION.** (A-B) DLS measurements of (A) hydrodynamic size and (B) PDI from P-SPION and MoNP-SPION; n = 3. (C) Representative transmission electron microscopy image of MoNP-SPION. Scale bar = 100 nm.

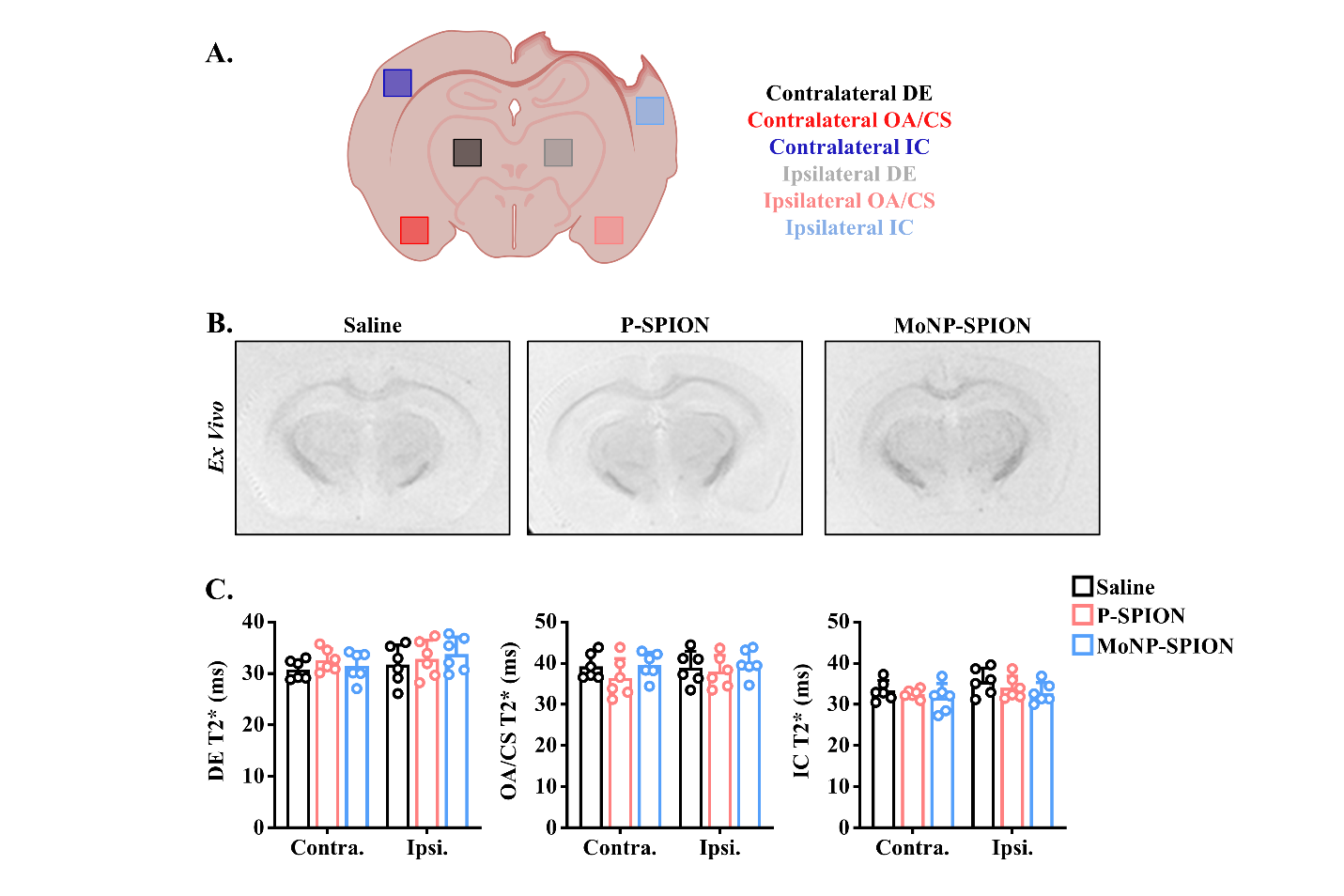

**Figure S5. *In vivo* MRI assessment in naïve mice.** (A) Schematic diagram of region-of-interest (ROI) placement. (B) Representative *in vivo* T2*-weighted coronal plane MR images of naïve mouse brains before and after administration of MoNP-SPION, P-SPION, and saline. (C) Quantification of post-injection T2* as a percentage of baseline values; n = 6. Data is presented as mean ± SD. Statistical significance was calculated with two-way ANOVA with Tukey's post-hoc multiple comparisons test.

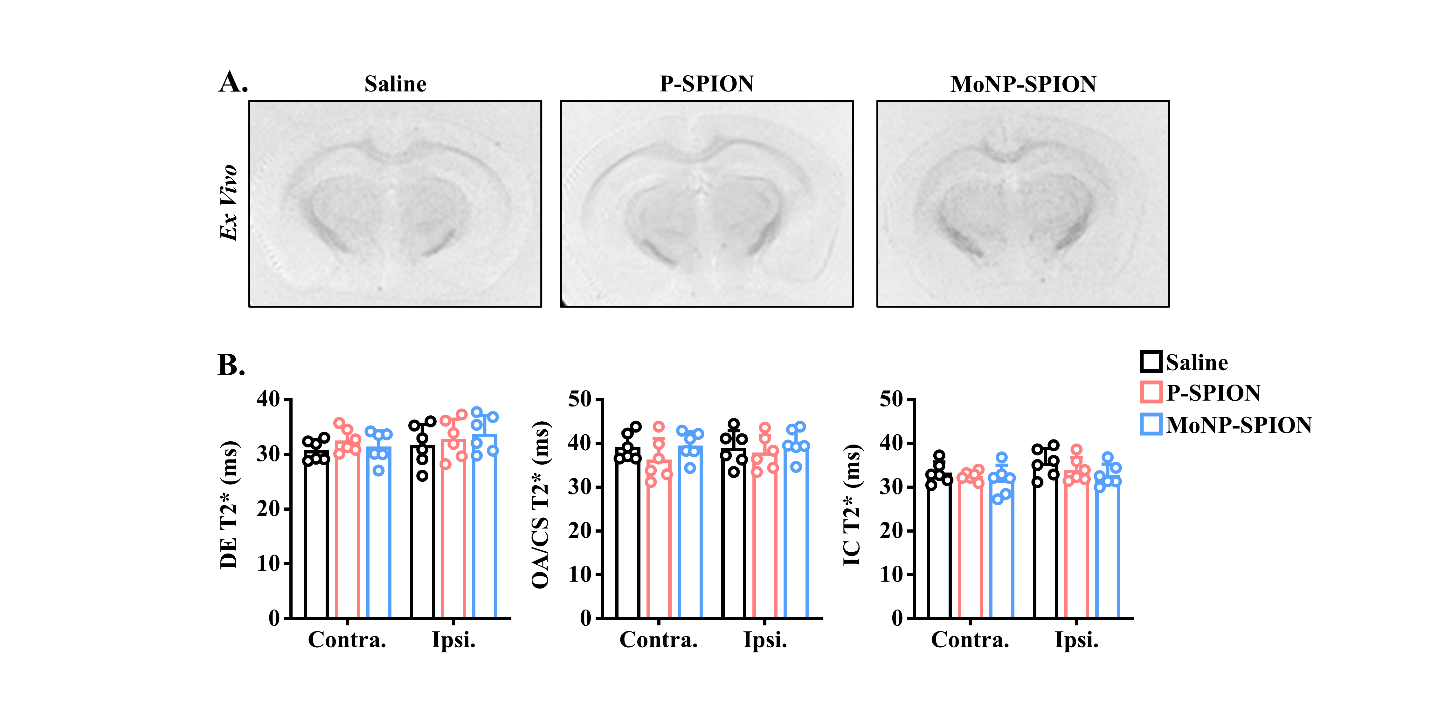

**Figure S6. *Ex vivo* MRI assessment in naïve mice.** (A) Representative *ex vivo* T2*-weighted images of naïve mouse brains after administration of MoNP-SPION, P-SPION, or saline. (B) Quantification of T2* within the DE (left), OA/CS (center), and IC (right) of the contralateral and ipsilateral hemispheres; n = 6. Data is presented as mean ± SD. Statistical significance was calculated using two-way ANOVA with Tukey's post-hoc multiple comparisons test.

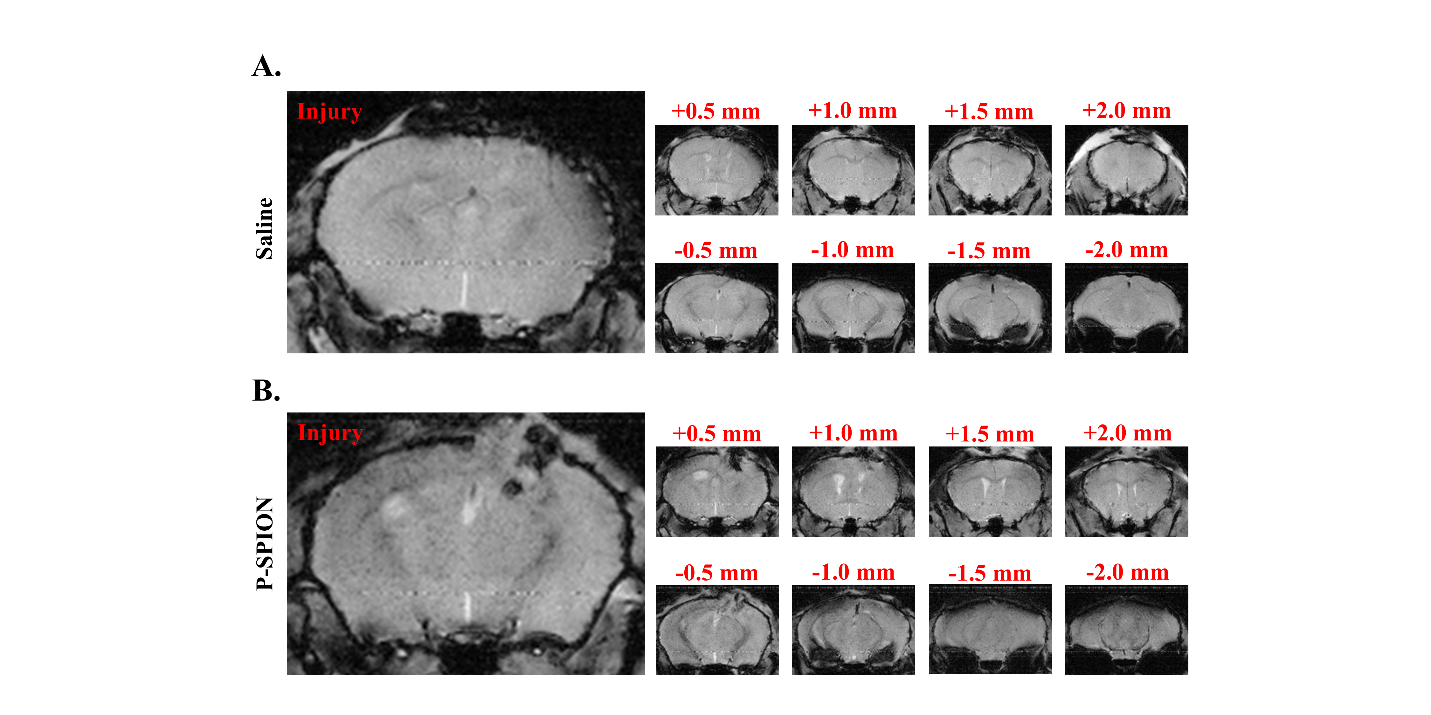

**Figure S7. *In vivo* multi-slice MRI of injured mouse brains following saline or P-SPION administration.** (A-B) Representative *in vivo* multi-slice coronal plane T2*-weighted MR images of mouse brains 1 day after injury, spanning from +2 mm rostral to -2 mm caudal of the injury core, following administration of (A) saline or (B) P-SPION.

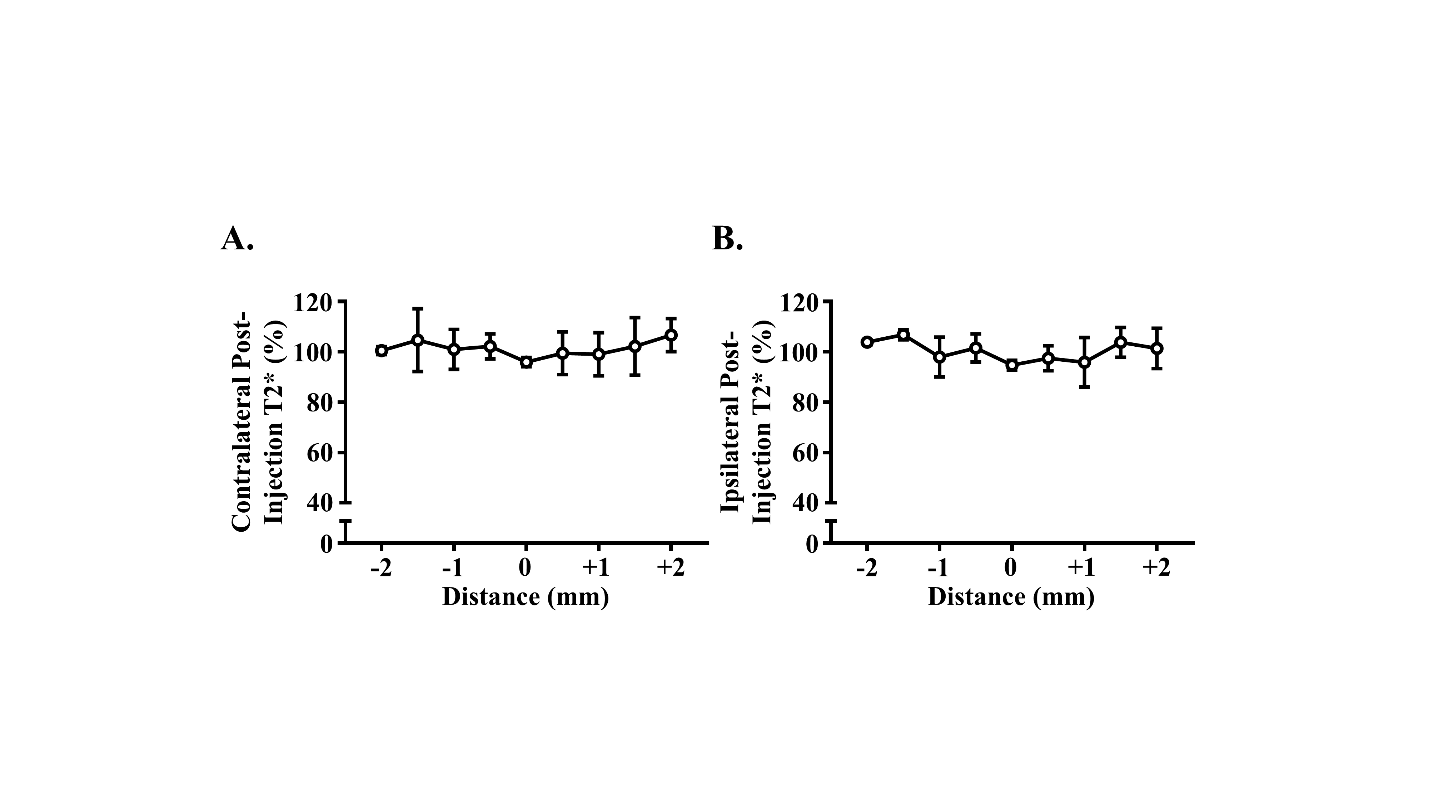

**Figure S8. Multi-slice T2* analysis of naïve mice following MoNP-SPION administration.** Post-injection T2* decay analysis of naïve mouse brain 3 hours after MoNP-SPION administration within the (A) contralateral and (B) ipsilateral hemispheres; n = 3. Data is presented as mean ± SD.

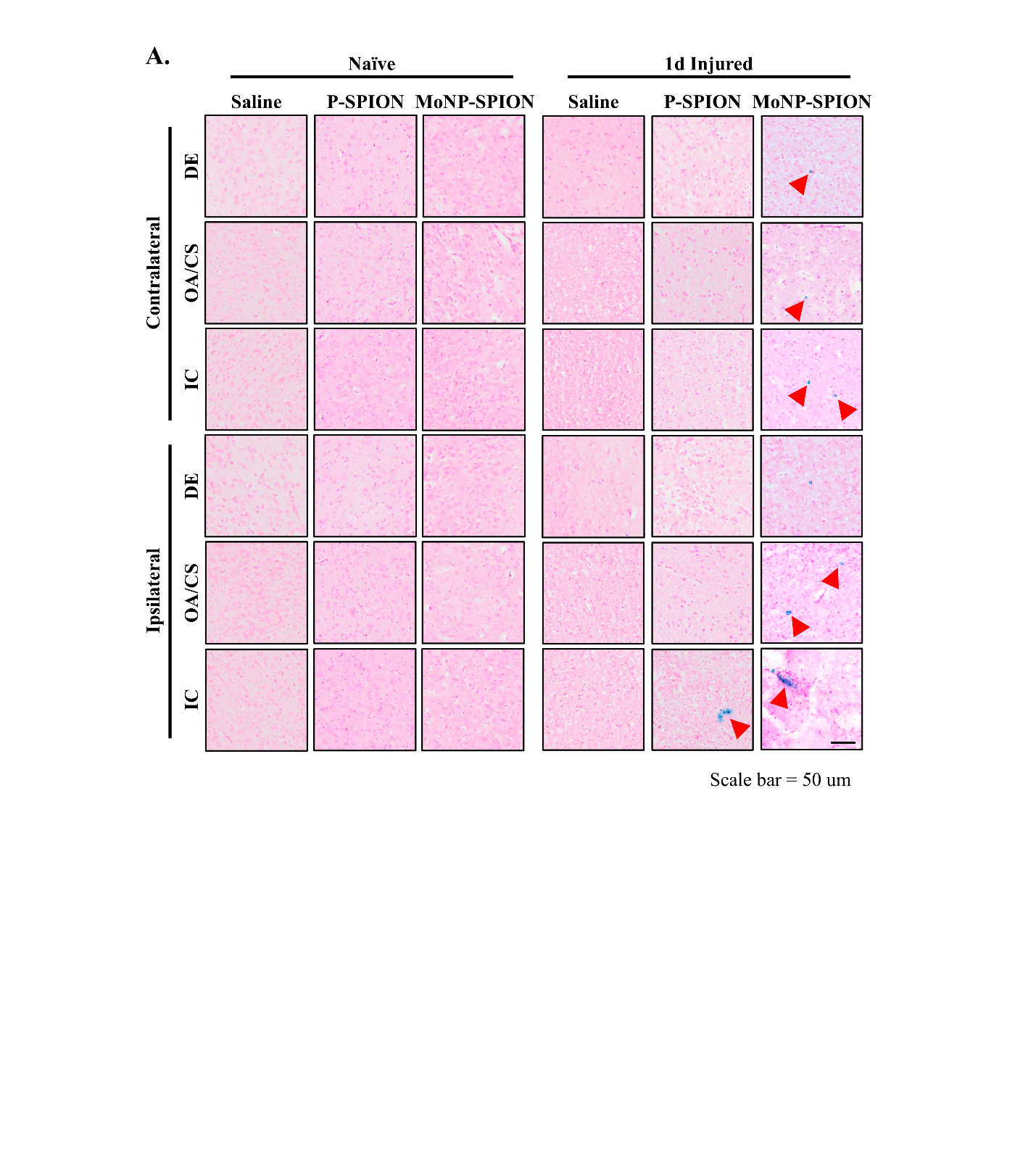

**Figure S9. Histology detection of iron deposition in injured mouse brains.** (A) Representative Prussian Blue and Nuclear Fast Red-stained brain sections from the DE, OA/CS, and IC of the contralateral and ipsilateral hemispheres following administration of MoNP-SPION, P-SPION, or saline. Red arrows indicate iron deposits. Scale bar = 50 µm.

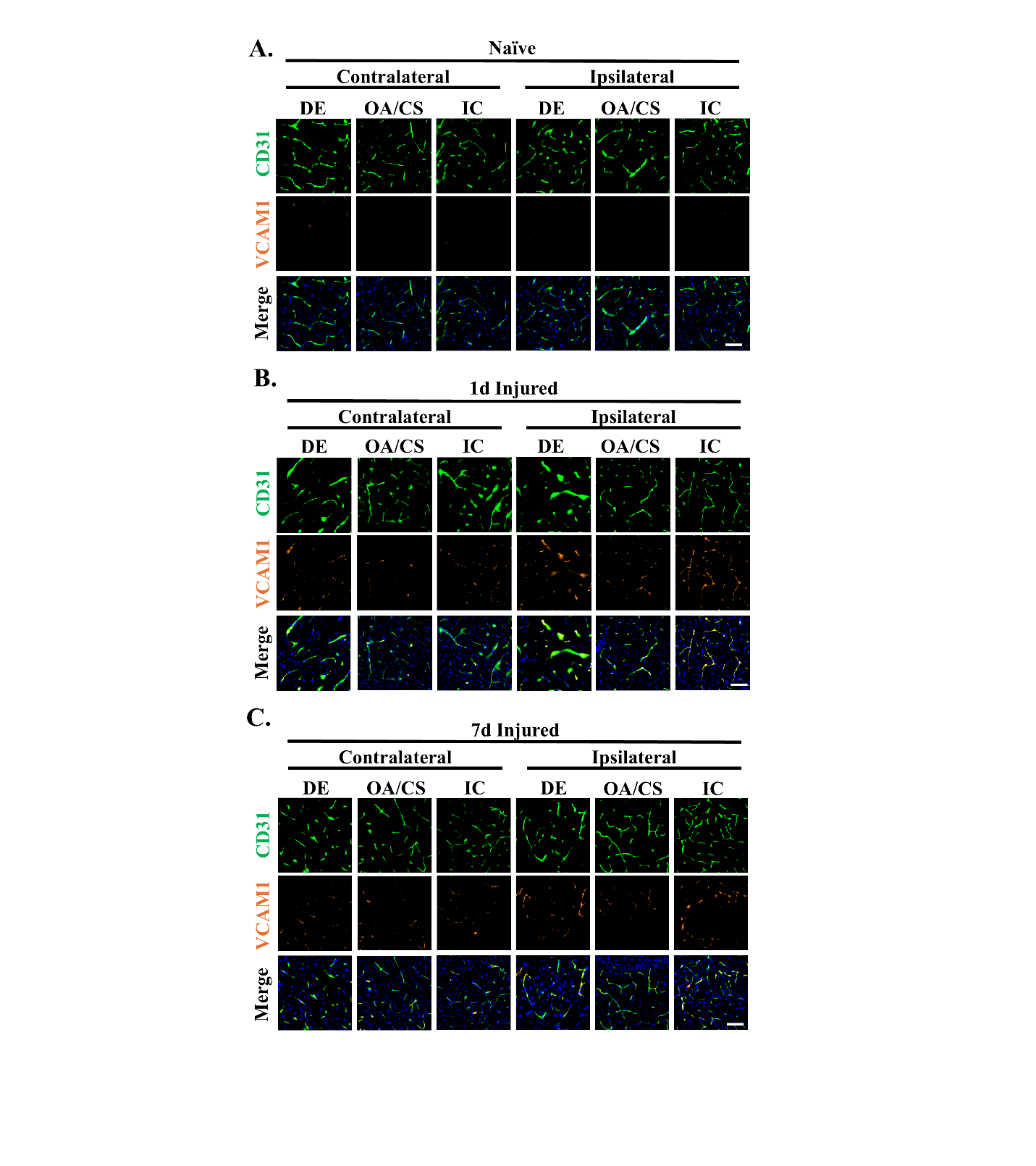

**Figure S10. Spatiotemporal VCAM1 expression after TBI.** (A) Schematic diagram of ROI placement. (B-D) CD31 (green) and VCAM1 (orange) staining and colocalization within the DE, OA/CS, and IC of the contralateral and ipsilateral hemispheres of (B) naïve, (C) 1-day injured, and (D) 7-day injured mouse brains. Scale bar = 50 µm.
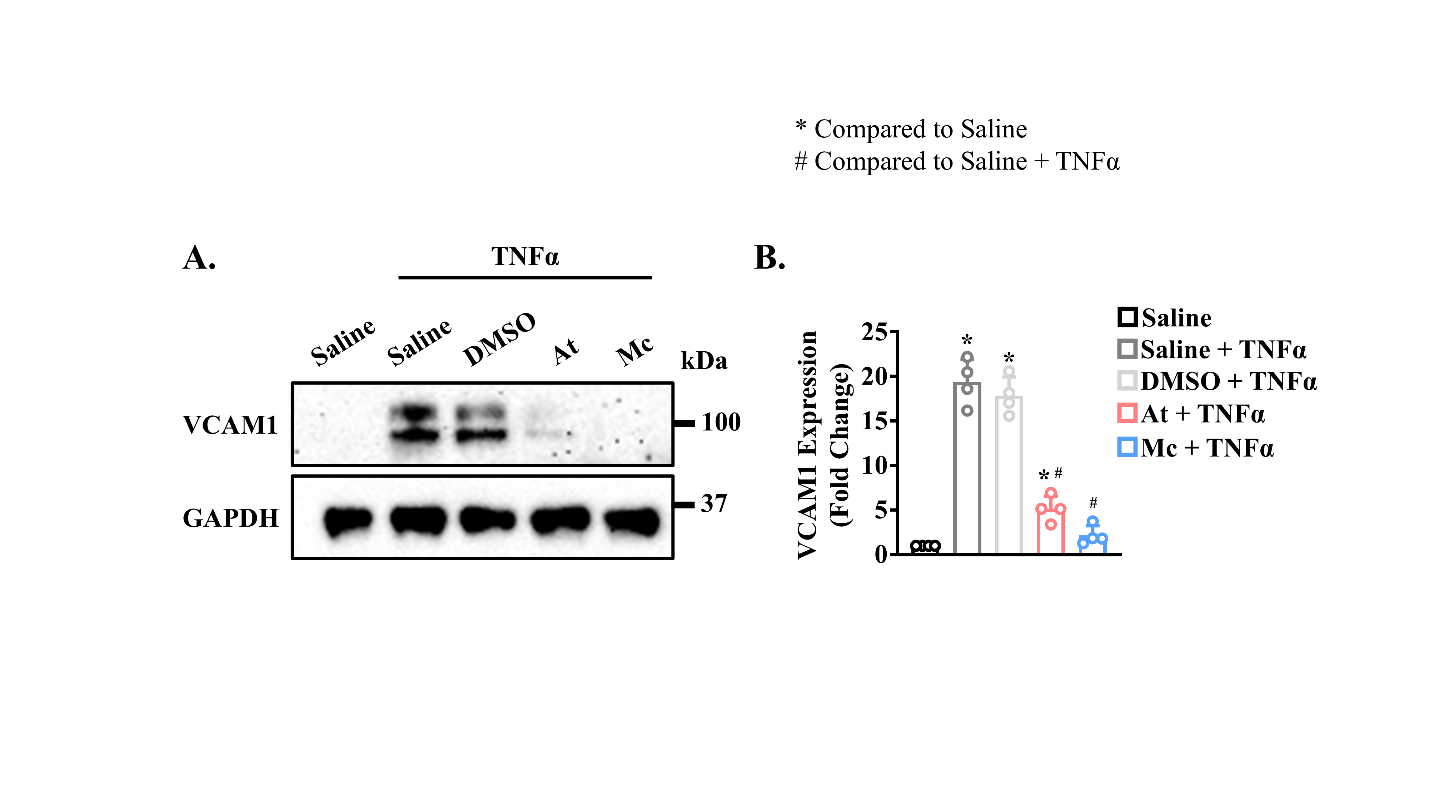

**Figure S11. Therapeutic suppression of VCAM1 suppression in TNFα-activated HBMECs.** (A) Representative Western blot image of VCAM1 and GAPDH from HBMECs pretreated with Mc, At, DMSO, and saline. (B) Quantification of VCAM1 expression; n = 4. Data is presented as mean ± SD. * indicates p < 0.05 compared to saline control; # indicates p < 0.05 compared to saline + TNFα. Statistical significance was calculated using one-way ANOVA with Tukey's multiple post-hoc comparisons test.

**SUPPLIMENTARY TABLES:**

|  |  |  | **Contralateral DE** | **Contralateral OA/CS** | **Contralateral IC** | **Ipsilateral DE** | **Ipsilateral OA/CS** | **Ipsilateral IC** |
| --- | --- | --- | --- | --- | --- | --- | --- | --- |
| **T2* (ms)** | **Naïve** | **Saline** | 30.79±1.90 | 39.28±3.15 | 33.36±2.44 | 31.76±3.80 | 38.98±3.99 | 35.61±3.24 |
|  |  | **P-SPION** | 32.59±2.22 | 36.42±4.77 | 32.60±1.07 | 32.90±3.60 | 37.97±3.93 | 34.06±2.80 |
|  |  | **MoNP-SPION** | 31.48±2.83 | 39.56±3.18 | 31.65±3.41 | 33.80±3.36 | 40.01±3.31 | 32.78±2.52 |
|  | **1d Injured** | **Saline** | 37.36±4.26 | 39.04±2.38 | 37.54±3.48 | 34.02±3.50 | 43.48±3.94 | 39.29±5.94 |
|  |  | **P-SPION** | 34.77±6.82 | 36.59±2.49 | 33.27±3.22 | 33.51±4.34 | 37.94±5.33 | 32.16±4.07 |
|  |  | **MoNP-SPION** | 29.08±3.86 | 31.52±2.29 | 26.29±6.15 | 24.26±6.90 | 29.78±2.91 | 24.01±5.70 |

**Table S1. *Ex vivo* T2* analysis in naïve and 1-day injured mice.** *Ex vivo* T2* decay constants acquired within the contralateral DE, contralateral OA/CS, contralateral IC, ipsilateral DE, ipsilateral OA/CS, and ipsilateral IC of naïve and 1 day injured mice after administration of MoNP-SPION, P-SPION, or saline. n = 6. Data is represented as mean ± SD.

|  |  | **Contralateral DE** | **Contralateral OA/CS** | **Contralateral IC** | **Ipsilateral DE** | **Ipsilateral OA/CS** | **Ipsilateral IC** |
| --- | --- | --- | --- | --- | --- | --- | --- |
| **T2* (ms)** | **Naïve** | 31.48±2.83 | 39.56±3.18 | 31.65±3.41 | 33.80±3.36 | 40.01±3.31 | 32.78±2.52 |
|  | **1d Injured** | 29.08±3.86 | 31.52±2.29 | 26.29±6.15 | 24.26±6.90 | 29.78±2.91 | 24.01±5.70 |
|  | **7d Injured** | 29.40±2.81 | 28.22±4.89 | 29.08±1.68 | 25.97±2.77 | 30.43±2.19 | 25.65±2.06 |

**Table S2.** **Spatiotemporal *ex vivo* T2* analysis.** *Ex vivo* T2* decay constants acquired within the contralateral DE, contralateral OA/CS, contralateral IC, ipsilateral DE, ipsilateral OA/CS, and ipsilateral IC of naïve, 1 day injured, and 7 day injured mice after administration of MoNP-SPION. n = 6. Data is represented as mean ± SD.

|  |  | **Contralateral DE** | **Contralateral OA/CS** | **Contralateral IC** | **Ipsilateral DE** | **Ipsilateral OA/CS** | **Ipsilateral IC** |
| --- | --- | --- | --- | --- | --- | --- | --- |
| **T2* (ms)** | **Vehicle Control** | 27.85±1.76 | 29.67±3.19 | 27.47±2.00 | 27.20±1.77 | 28.90±2.14 | 25.42±1.56 |
|  | **At** | 29.82±1.96 | 30.17±3.03 | 29.68±2.91 | 27.95±1.03 | 29.53±1.86 | 27.20±2.08 |
|  | **Mc** | 33.62±2.51 | 31.98±3.66 | 33.68 ± 4.09 | 32.52±2.78 | 34.27±2.68 | 29.93±1.57 |

**Table S3. *Ex vivo* T2* analysis after 7-day anti-inflammatory treatment.** *Ex vivo* T2* decay constants acquired within the contralateral DE, contralateral OA/CS, contralateral IC, ipsilateral DE, ipsilateral OA/CS, and ipsilateral IC after administration of MoNP-SPION in injured mice administered seven days of Mc, At, or a vehicle control treatment. n = 6. Data is represented as mean ± SD.
